# Integrated genomic analysis of AgRP neurons reveals that IRF3 regulates leptin’s hunger-suppressing effects

**DOI:** 10.1101/2022.01.03.474708

**Authors:** Frankie D. Heyward, Nan Liu, Christopher Jacobs, Rachael Ivison, Natalia Machado, Aykut Uner, Harini Srinivasan, Suraj J. Patel, Anton Gulko, Tyler Sermersheim, Stuart H. Orkin, Linus Tsai, Evan D. Rosen

**Affiliations:** Division of Endocrinology, Diabetes, and Metabolism, Beth Israel Deaconess Medical Center; Cancer and Blood Disorders Center, Dana-Farber Cancer Institute and Boston Children’s Hospital; Howard Hughes Medical Institute, Boston, MA, USA; Bone Marrow Transplantation Center of the First Affiliated Hospital, Zhejiang University School of Medicine, 310003 Hangzhou, China; Liangzhu Laboratory, Zhejiang University Medical Center, Hangzhou, 311121, China; Department of Neurology, Beth Israel Deaconess Medical Center; Division of Gastroenterology & Hepatology, Beth Israel Deaconess Medical Center; Broad Institute of Harvard and MIT; Harvard Medical School

## Abstract

AgRP neurons in the arcuate nucleus of the hypothalamus (ARC) coordinate homeostatic changes in appetite associated with fluctuations in food availability and leptin signaling. Identifying the relevant transcriptional regulatory pathways in these neurons has been a priority, yet such attempts have been stymied due to their low abundance and the rich cellular diversity of the ARC. Here we generated AgRP neuron-specific transcriptomic and chromatin accessibility profiles during opposing states of fasting-induced hunger and leptin-induced hunger suppression. Cis-regulatory analysis of these integrated datasets enabled the identification of 28 putative hunger-promoting and 29 putative hunger-suppressing transcriptional regulators in AgRP neurons, 16 of which were predicted to be transcriptional effectors of leptin. Within our dataset, Interferon regulatory factor 3 (IRF3) emerged as a leading candidate mediator of leptin-induced hunger-suppression. Gain- and loss-of-function experiments *in vivo* confirm the role of IRF3 in mediating the acute satiety-evoking effects of leptin in AgRP neurons, while live-cell imaging *in vitro* indicate that leptin can activate neuronal IRF3 in a cell autonomous manner. Finally, we employ CUT&RUN to uncover direct transcriptional targets of IRF3 in AgRP neurons *in vivo*. Thus, our findings identify AgRP neuron-expressed IRF3 as a key transcriptional effector of the hunger-suppressing effects of leptin.

## INTRODUCTION

Increasing the granularity with which we understand the homeostatic regulation of energy balance (i.e., food intake and energy expenditure) will represent a major step towards the ultimate goal of treating obesity. It has long been recognized that the arcuate nucleus (ARC) of the hypothalamus is indispensable for maintaining energy homeostasis (Andermann and Lowell, 2017). Amid the rich diversity of cell types within the ARC, AgRP neurons play a critical role in coordinating the physiological processes needed to maintain energy homeostasis in the face of changing energy availability.

AgRP neurons are activated during states of negative energy balance (i.e., fasting) and selective chemogenetic and optogenetic activation of these neurons drives mice to become markedly hyperphagic (Andermann and Lowell, 2017). Conversely, post-developmental genetic deletion of AgRP neurons renders mice profoundly hypophagic, and during periods of simulated energy repletion, the satiety-evoking adipokine leptin has been shown to decrease the firing rate of AgRP neurons (Beutler et al., 2017; Luquet et al., 2005). Finally, deletion of the leptin receptor specifically from adult mouse AgRP neurons results in mice that phenocopy the morbid obesity exhibited by *db/db* whole-body leptin receptor null mice, indicating that leptin’s anti-obesity effects can be mediated via its direct effects on AgRP neurons (Xu et al., 2018). Given these findings, much effort has been directed towards elucidating the transcriptional programs that allow AgRP neurons to respond appropriately to changes in energy availability.

Distinct alterations in the transcriptional profiles of AgRP neurons have been noted in response to fasting, including genes encoding neurotransmitter receptors, ion channels and secreted proteins (Henry et al., 2015). These transcriptional changes are suspected to underlie the requisite cell type-specific alterations in neuronal excitability that enable AgRP neurons to influence downstream satiety signaling circuits.

Transcriptional regulation is a key mechanism by which leptin confers its effects on neurons. Mice with a pan-neuronal deletion of signal transducer and activator of transcription 3 (STAT3), the best-known transcriptional effector of leptin action, exhibit profound obesity which is similar to that seen in leptin receptor null mice (Bates et al., 2003; Gao et al., 2004; Hummel et al., 1966). Moreover, mice devoid of STAT3 specifically in AgRP neurons exhibit an increase in body weight compared to control mice (Gong et al., 2008). Activating transcription factor 3 has also been shown to mediate some of the effects of leptin *in vivo* (Allison et al., 2018). Additional evidence of a transcriptional basis for leptin comes from the observation that leptin’s suppressive effects on AgRP neuronal activity occur gradually over a 3-hour time course, a finding that implicates the involvement of transcriptional regulation (Beutler et al., 2017).

Previous studies have characterized the transcriptional events that occur in all leptin-receptor expressing cells in the hypothalamus in response to leptin (Allison et al., 2018). More recently, one group assessed AgRP neuron specific transcriptional changes in response to fasting and leptin-administration, observing 33 leptin-induced transcriptional changes that were linked to biological processes suspected to facilitate behavioral changes in response to fluctuations in energy availability (Cedernaes et al., 2019). Another group generated a chromatin accessibility landscape of the broad population of leptin receptor-expressing cell types in the brain, yet no bioinformatic assessment of leptin-induced changes in the epigenetic landscape was offered (Inoue et al., 2019). To date, we lack a rich and comprehensive understanding of the transcriptional and transcriptional regulatory changes that occur in AgRP neurons in response to leptin.

High throughput DNA sequencing techniques that generate genome-wide profiles of open chromatin regions (OCRs) have been leveraged to infer the identity of transcription factors (TFs) that govern the cellular identity and activity state of a variety of cell-types (Guan et al., 2020; Hiraike et al., 2017; Mikkelsen et al., 2010; Stroud et al., 2020; Su et al., 2017). However, applying this approach to study transcriptional regulation within purified populations of AgRP neurons has been challenging for two reasons. The first has to do with the rich cellular heterogeneity within the ARC, where over 50 cell-types have been identified, which necessitates the isolation of these cells prior to their being studied (Campbell et al., 2017). The second is owed to the relatively low abundance of mouse AgRP neurons (∼9,000 per mouse) (Betley et al., 2013) compared to the cellular input typically required for traditional deeply sequenced genomic profiling techniques (≥ 100,000). To surmount these two obstacles, we developed mouse lines in which AgRP neurons express tagged ribosomes and nuclei, thereby enabling the generation of transcriptomic and epigenomic profiles from as few as 10,000 cells (Roh et al., 2018). This enabled us to assess the transcriptomic and cis-regulatory element landscape of AgRP neurons during opposing states of energy availability as a means of identifying additional transcriptional pathways that underlie the distinct cellular states exhibited by these neurons during periods of hunger and satiety.

## RESULTS

### Validating a transgenic tool for obtaining AgRP neuron-specific RNA-seq profiles

We previously developed a mouse model capable of providing cell type-specific transcriptional and epigenomic profiles from complex tissues, which we call NuTRAP (Nuclear tagging and Translating Ribosome Affinity Purification) (Roh et al., 2018). NuTRAP mice possess a loxP-stop-loxP sequence immediately upstream of a cassette that expresses a GFP-tagged L10a ribosomal subunit as well as mCherry-tagged RANGAP, which localizes to the nuclear membrane. Nuclei can be isolated by flow-sorting for either mCherry or GFP, as the GFP-tagged L10a ribosomal subunit is enriched in the nucleolus, thereby allowing for nuclear sorting. Crossing NuTRAP mice with either AgRP-IRES-Cre or POMC-IRES-Cre mice generates mice with tagged ribosomes or nuclei selectively within AgRP or POMC neurons of the ARC (here referred to as NuTRAP^AgRP^ or NuTRAP^Pomc^). Using this approach, tagged ribosomes can be isolated via affinity purification for RNA-seq, and tagged nuclei can be isolated via fluorescence-activated nuclear sorting (FANS) for ATAC-seq. We confirmed the viability of using NuTRAP^AgRP^ mice to generate cell-type specific transcriptional profiles by observing distinct gene expression patterns for GFP+ (AgRP neuron positive) and whole sample input ribosomes isolated from NuTRAP^AgRP^ mice using qPCR. Canonical AgRP neuron genes were enriched, including *Agrp* and *Npy*, whereas genes known to be de-enriched in AgRP neurons were found at reduced levels, such as *Pomc*, *Gfap*, *Mobp* (**Figure S1A**). We also observed that TRAP performed on four pooled ARCs resulted in greater enrichment for AgRP neuron markers than that seen with a smaller number of ARCs (**Figure S1A**). To overcome the observed minimal input requirements for, and the reliability of, TRAP we opted to use four pooled ARCs for the remainder of our TRAP studies. Next, RNA-seq was used to validate and extend our qPCR data. Again, as anticipated, the canonical marker genes *AgRP* and *Npy* were enriched in our AgRP neuronal population (**Figure S1B**). We also observed enrichment for other genes known to be expressed in these neurons, including *Corin*, *Otp, Ghsr,* and *Acvr1c* (Henry et al., 2017). Notably, the *Serpina3* family of genes was markedly enriched in AgRP neurons, including *Serpina3c,* the third-most enriched transcript, along with *Serpina3i*, *Serpina3n, Serpina3g*, and *Serpina3h*, suggesting that this family might play an important role in the regulation AgRP neuron biology. *Lepr*, encoding the leptin receptor, was also significantly enriched in AgRP neurons. We then compared the transcriptomic profiles of NuTRAP^AgRP^ and NuTRAP^Pomc^ GFP+ TRAP samples. POMC neuron marker genes *Pomc* and *Cartpt* were among the most POMC neuron-enriched genes, along with *Prdm12*, which is indispensable for the expression of *Pomc* (**Figure S1C**)(Hael et al., 2020). Based on these results we were confident that we could isolate and transcriptionally profile a purified population of AgRP neurons.

### Obtaining AgRP neuron-specific transcriptional profiles in response to fasting and leptin

We next sought to assess transcriptional profiles of AgRP neurons during various states of energy balance using NuTRAP^AgRP^ mice that were fed, fasted and leptin-treated (n = 3-4 pooled samples per experimental condition, 4 pooled mouse ARCs per sample). Mice were either provided food *ad libitum* or were fasted overnight and then injected with either (5mg/kg) i.p. leptin or vehicle; three hours later mice were sacrificed and the ARC dissected out for further study (**Figure 1A**). The dose of leptin chosen was confirmed to activate leptin-signaling-mediated activation of STAT3 (assessed by phospho-STAT3) in the ARC, with substantial co-localization within the cell body of AgRP neurons (**Figure 1B**). We identified 527 and 295 genes that were induced and repressed by fasting in AgRP neurons, respectively (**Figure 1C**). Among the fasting-induced genes were *AgRP*, *Npy*, and *Vgf*, all shown previously to be affected by food withdrawal (Campbell et al., 2017; Henry et al., 2015). Other genes exhibiting increased expression with fasting include *Acvr1c*, *Lyve1*, *Svs2* (**Figure 1D**). WebGestalt (WEB-based Gene SeT AnaLysis Toolkit) gene set enrichment analysis (GSEA) for geneontology (GO) biological process revealed fasting-induced transcriptional programs that were associated with various molecular functions including, “response to endoplasmic reticulum stress” and “cytoplasmic translation”, and “ribonucleoprotein complex biogenesis”, among others (**Figure 1E**). These findings recapitulated our earlier work, and that of another group (Campbell et al., 2017; Henry et al., 2015). Within our dataset exist fasting-induced genes that are categorized as ion channels (e.g., *Kcnv2*, *P2rx5*, *Cftr*), G-protein coupled receptors (e.g., *Adgrg2*, *Gprc5a*, *Gpr6*), secreted proteins (e.g., *Agrp*, *Svs2*, *Vgf)*, transporters (e.g., *Rbp4*, *Sec61b*, *Sc2c*), and transcriptional regulators (e.g., *Sox4*, *Nfil3*, and *Npas4*; **Figure 1G**). Unsurprisingly, many transcripts encoding classic immediate early genes (IEGs) were upregulated in AgRP neurons during fasting (e.g., *Nr4a1*, *Fos*, *Myc*, *Fosb*, *Egr1*, *Junb*), potentially driven by the sustained increase in neuronal firing that occurs throughout the course of food deprivation (Takahashi and Cone, 2005).

**Figure 1.**
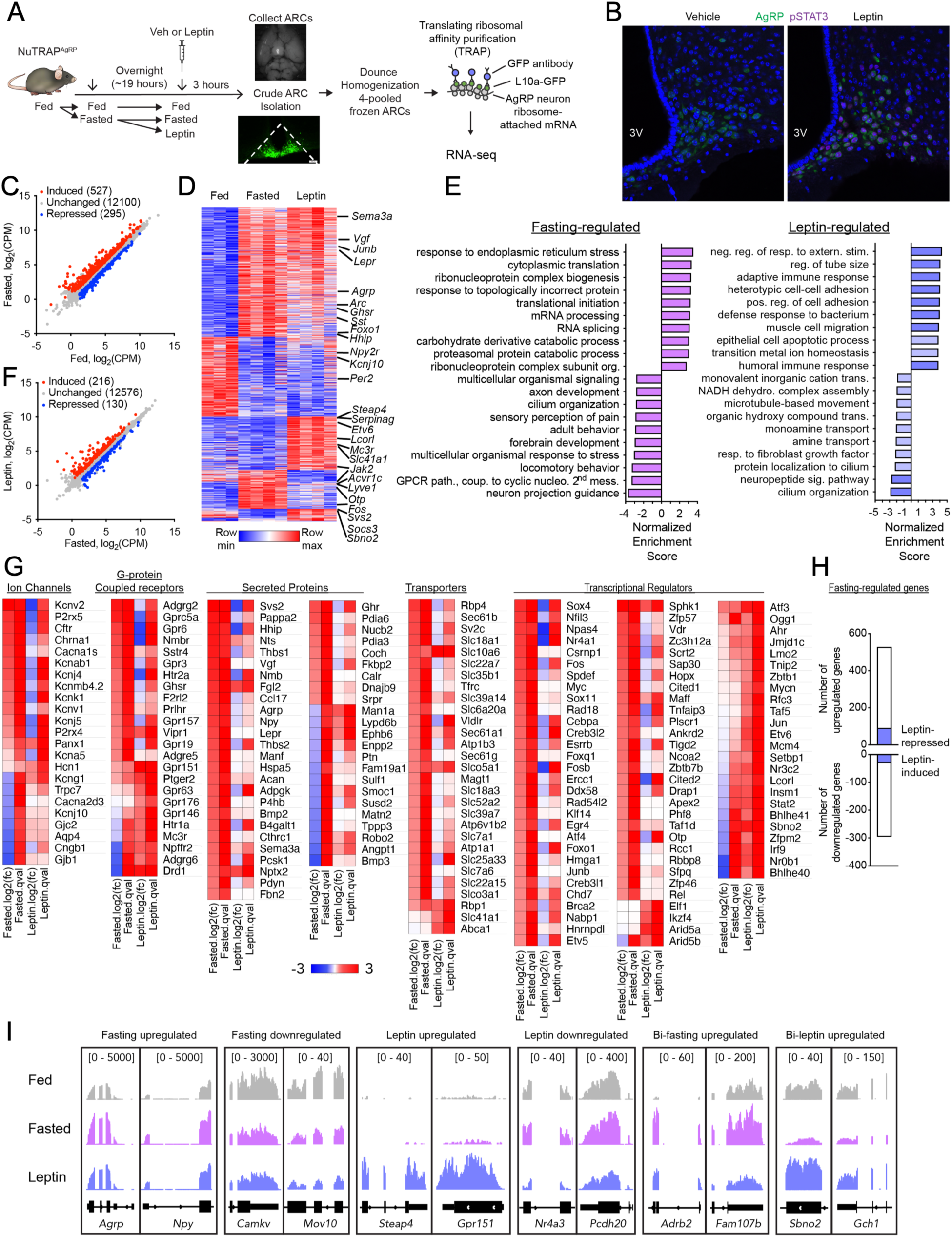
Transcriptomic profiles of AgRP neurons during opposing states of energy balance. (**A**) Schematic of the experimental design. NuTRAP^AgRP^ mice were fed, then a subset were fasted overnight (∼19hr), following which mice were treated with IP vehicle or leptin (5mg/kg). After 3 hours, mice were euthanized and ARCs were isolated, frozen, and homogenized according to the TRAP protocol. (**B**) Representative immunofluorescence image of ARC revealing induction of pSTAT3 in response to leptin administration (right) compared to vehicle treatment (left). (**C**) Scatter plot showing comparing expression of genes within AgRP neurons from fasted versus fed littermates (*n* = 4, 3) [fold change >0.5 up (red) or down (blue), false discovery rate (FDR) < 0.05]; CPM, counts per million. The number of genes in a given expression profile is in parentheses. (**D**) Heatmap of differentially expressed genes in fed, fasted, and leptin-treated mice. (**E**) WebGestalt (WEB-based Gene SeT AnaLysis Toolkit) gene set enrichment analysis (GSEA) for geneontology biological process in either fasting-regulated (left, induced and repressed), or leptin-regulated (right, induced and repressed) transcriptional pattern categories with Normalized Enrichment Scores. All bars have a FDR < 0.05 except those that are light blue in the leptin-regulated condition (**F**) Scatter plot showing regulated genes within AgRP neurons from leptin-treated versus fasted littermates (*n* = 4, 4) (fold change > 0.5 up (red) or down (blue), FDR < 0.05). CPM, counts per million. The number of genes in a given expression profile is in parentheses. (**G**) Heatmaps of selected gene categories. (**H**) Top: bar graphs showing the number of fasted up-regulated genes (N = 527) that are also down-regulated with leptin (blue bars, N = 90). Bottom: bar graphs showing the number of fasted down-regulated genes (N = 295) that are also up-regulated with leptin (blue bars, N = 30). (**I**) Genome browser views (in Integrative Genomics Viewer (IGV)) of RNA-seq tracks of representative fasting-upregulated, fasting-downregulated, leptin-upregulated, leptin-downregulated, bidirectional fasting upregulated (Bi-fasting upregulated), and bidirectional leptin upregulated (Bi-leptin upregulated) genes.

In leptin-treated mice, we observed the induction of 216, and repression of 130, genes in AgRP neurons (**Figure 1F**). *Socs3*, a component of the leptin signaling negative feedback loop, was significantly upregulated in response to leptin-treatment. Notably, leptin-induced transcriptional programs were associated with biological processes including “negative regulation of response to external stimulus”, “regulation of tube size”, and “adaptive immune response”, among others (**Figure 1E**). Leptin-induced genes included those encoding ion channels (e.g., *Trpv6*, *Kcng1*), G-protein coupled receptors (e.g., *Drd1*, *Adgrg6*, *Gpr151*), secreted proteins (e.g., *Man1a*, Lypd6b, Ephb6), transporters (e.g., *Rbp4*, *Slc41a1*, *Abca1*), and transcriptional regulators (e.g., *Stat2*, *Bhlhe41*, and *Sbno2*)(**Figure 1G**).

We were particularly interested in identifying “bidirectional” transcriptional changes in response to fasting and leptin (i.e., genes induced by fasting and repressed by leptin, or vice versa). This analysis revealed 120 genes exhibiting a bidirectional expression pattern, including 90 genes induced by fasting and repressed by leptin (e.g., *Acvr1c*, *Gpr157*, *Lyve1*, and *Otp)*, and 30 repressed by fasting and induced in response to leptin (e.g., *Sbno2*, *Serpina3i*, *Drd1*, *Man2a1*) (**Figure 1H, 1I**). Expression of this set of 30 genes is reduced in response to fasting, a state that has classically been associated with a marked decline in serum leptin levels and leptin signaling, while their expression is increased in response to the administration of exogenous leptin in the fasted state (Ahima et al., 1996). Given the apparent bi-directional leptin sensitivity of these 30 genes, it is conceivable that leptin depletion-driven declines in the expression of these genes may mediate the adaptive physiological response to starvation attributed to the loss of leptin in the fasted state (Ahima et al., 1996). We also performed TRAP-seq on fed, fasted and leptin-treated NuTRAP^Pomc^ samples and observed relatively few significantly changed genes in response to fasting, a finding that recapitulated earlier work, and prompted us to focus on AgRP neurons for the remainder of our studies (**Figure S1D**)(Henry et al., 2015).

### Validating a transgenic tool for obtaining AgRP neuron-specific ATAC-seq profiles

Next, we confirmed the feasibility of using NuTRAP mice to assess chromatin state in AgRP neurons. Using pooled NuTRAP^AgRP^ mouse ARCs, we isolated and sorted 10,000 nuclei, and then performed Assay for Transposase Accessible Chromatin with high-throughput sequencing (ATAC-seq) for AgRP positive and negative populations (**Figures 2A, 2B**). We detected 15,530 and 9,029 closed chromatin regions that are specifically enriched in AgRP positive neurons (**Figure S2A**), and principal components analysis (PCA) showed a clear distinction between the two populations in terms of their respective chromatin states (**Figure S2B**). We next examined the chromatin state of AgRP neurons at key genes expressed in AgRP neurons, compared to that of the non-AgRP neuron population, and observed that AgRP neurons exhibited a selective enrichment of open chromatin regions (OCRs) at the promoters and enhancers of *Agrp*, *Npy*, and *Corin*, among others (**Figure S2C**). We also observed multiple OCRs upstream of *Lepr* in AgRP neurons (**Figure S2D**). Interestingly, these *Lepr*-associated OCRs are distinct from those observed in adipocytes, suggesting that putative enhancer elements for *Lepr* may be developmentally divergent (data not shown). We also detected OCR de-enrichment at the promoter of *Slc17a6*, which encodes the protein vesicular glutamate transporter 2 (VGlut2), reflecting the fact that AgRP neurons are gabaergic, rather than glutamatergic (**Figure S2C**). Moreover, de-enrichment of OCRs was noted at non-neuronal marker genes, such as those for oligodendrocytes (e.g., *Olig1* and *Olig2*) and tanycytes (e.g., *Sox2*), supporting the utility and fidelity of our method for obtaining low-input AgRP neuron-specific ATAC-seq profiles (**Figure S2C**).

**Figure 2.**
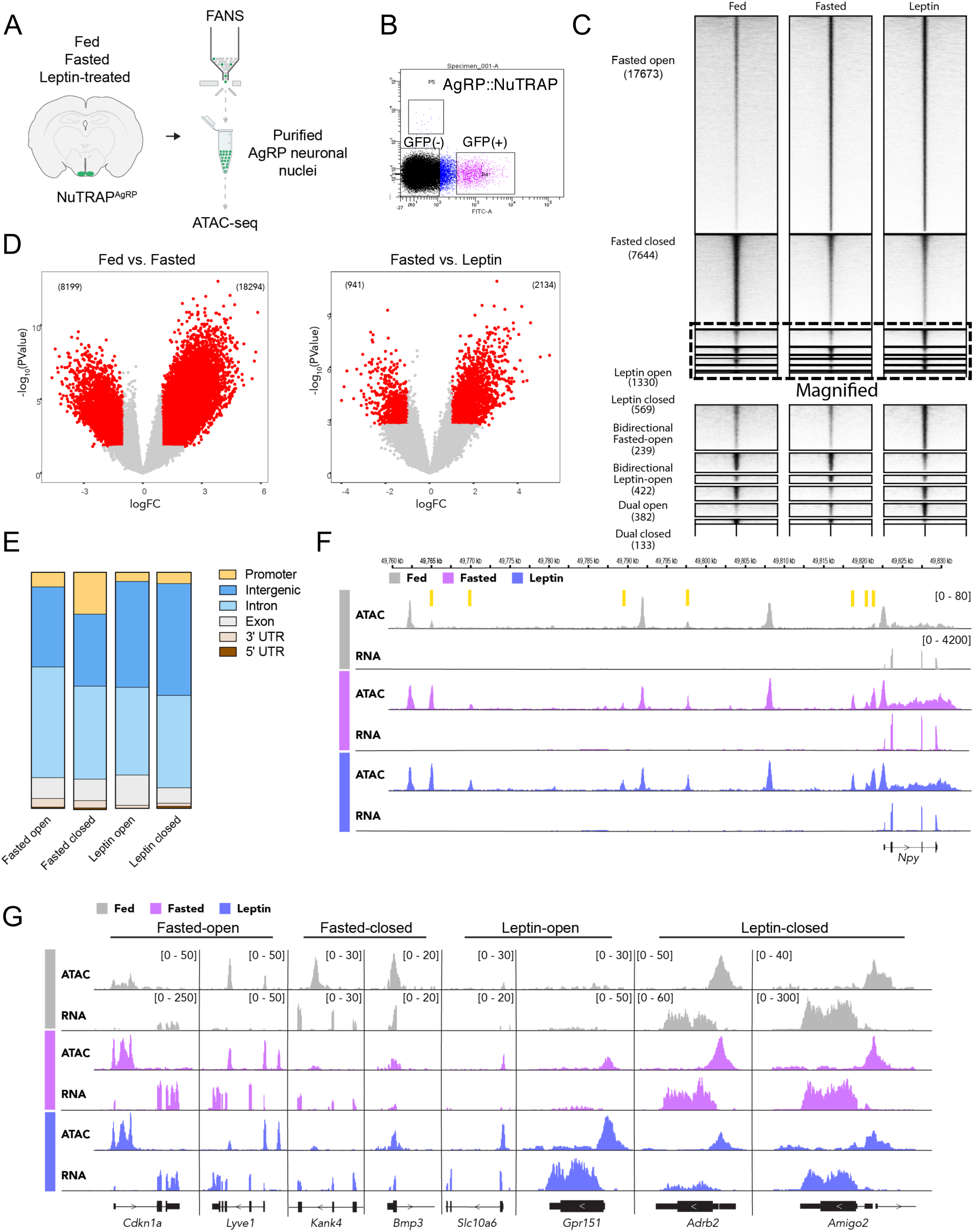
Alterations in energy availability impact the chromatin landscape of AgRP neurons. (**A**) Experimental schematic. ARCs from fed, fasted, and leptin-treated NuTRAP^AgRP^ mice were isolated followed by nuclear isolation. 10,648 ± 1489 nuclei, from 4 mice, were pooled for each condition (fed, fasted, leptin-treated) before being subjected to ATAC-seq. (**B**) FACS plot of single nuclei with FITC (GFP) fluorescence on the x-axis and mCherry fluorescence on the y-axis, with filtering gates showing discrete populations for NuTRAP^AgRP^ GFP+ and GFP-nuclei. (**C**) Heatmap showing differentially expressed OCRs for each of the treatment conditions clustered into one of eight patterns (rows). Amplitude of each peak center (±5 kb) is represented in black as indicated. (**D**) Volcano plots showing differentially expressed OCRs in fasted vs. fed mice (left, n = 2, 2) or leptin vs. fasted mice (right, n = 2, 2). Red dots correspond to significantly different gained-open and gained-closed regions [fold change >0.5 up (red) or down (red), FDR < 0.05]; CPM, counts per million. (**E**) Bar graphs revealing the genomic features of fasted-opened, fasted-closed, leptin-opened, leptin-closed ATAC-seq peaks. (**F**) Genome browser views (IGV) ofATAC-seq peaks within the vicinity of the *Npy* gene. Yellow bars correspond to fasted-opened ATAC-peaks. (**G**) Genome browser views (IGV) of representative Fasted-opened, Fasted-closed, Leptin-opened, and Leptin-closed ATAC-seq peaks near the indicated genes.

### AgRP neuron-specific ATAC-seq profiles in response to fasting and leptin

We next performed ATAC-seq on AgRP neurons from fed, fasted, and leptin-treated NuTRAP^AgRP^ mice. Of the 90,196 called ATAC-seq peaks in our dataset, we detected 18,294 fasted-opened and 8,189 fasted-closed regions, and 2,134 leptin-opened and 941 leptin-closed regions of chromatin (**Figures 2C, 2D**). We also detected changes in chromatin accessibility that were bidirectional (i.e., reciprocally regulated in fasted vs. leptin-treated) or dually enriched (i.e., open in response to both fasting and leptin-treatment) (**Figure 2C**). The majority of differentially expressed ATAC peaks were found within intergenic (36-45%) and intronic (43-50%) regions, while only 3-11% of ATAC-peaks were found within gene promoter regions, defined as regions <1000 bps upstream of the transcriptional start site (TSS) (**Figure 2E**), a changed genomic distribution of changed chromatin accessibility previously exhibited by activated neurons (Su et al., 2017). We observed areas of chromatin that became more accessible with fasting (i.e., fasted-opened) near various genes known to be enriched in response to fasting in AgRP neurons, including *Npy* (**Figures 2F, 2G**). Moreover, we also observed areas of chromatin that became less accessible with fasting (*i.e*., fasted-closed), as well as those that became more or less accessible with leptin (*i.e*., leptin-opened and leptin-closed) (**Figure 2G**).

### Uncovering transcriptional regulators that control AgRP neuron biology

Having generated an extensive atlas of dynamic chromatin regions in AgRP neurons in response to fasting and leptin treatment, we next sought to determine which TFs might bind to these regions. To accomplish this we performed Analysis of Motif Enrichment (AME), a program of the MEME suite, on chromatin regions with a high likelihood of impacting fasting- and leptin-induced transcriptional changes (**Figure 3A**) (Bailey et al., 2015). To accomplish this, we first filtered ATAC-seq peaks to retain those within ± 200 kb of a TSS corresponding to a gene whose expression is concordantly regulated (e.g., fasting-induced, gained-open ATAC-seq peaks near fasting-induced genes; hereafter called concordant fasted-opened; **Figures 3A, S3A**). We observed that 96%, 78%, 59%, and 51% of fasting-induced, fasting-repressed, leptin-induced, and leptin-repressed genes, respectively, were associated with a concordant peak (**Figure S3B**). The majority of genes associated with a concordant peak were in fact associated with numerous concordant peaks; most genes were associated with 4-6 peaks for fasting-induced genes, and 1-3 peaks per genes for the other three comparisons (**Figure S3C**). We also observed a positive relationship between the number of concordant peaks and the fold-change of the associated gene (**Figure S3D**). The number of concordant peaks identified for each of the four comparisons was 2452 fasted-opened, 531 fasted-closed, 203 leptin-opened, and 74 leptin-closed (**Figure 3B**).

**Figure 3.**
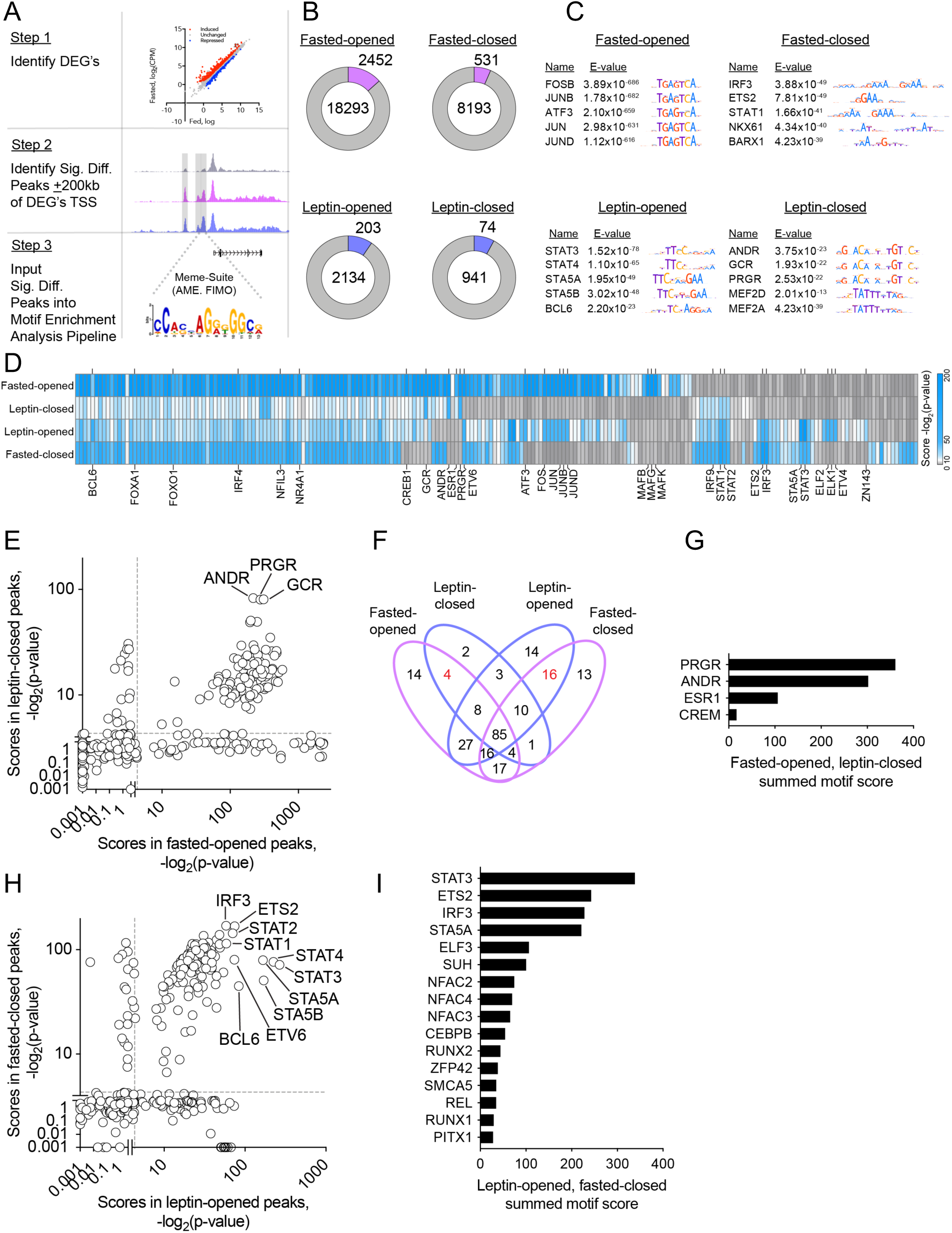
Integrated transcriptomic and cistromic analysis identifies putative leptin-sensitive TFs in AgRP neurons. (**A**) Schematic showing the broad computational heuristic employed to identify putative pro-satiety TF motifs. Step 1: Identify fasted-induced, fasting-repressed, leptin-induced, and leptin-repressed differentially expressed genes (DEGs). Step 2: Determine significantly different fasted-opened, fasted-closed, leptin-opened, and leptin-closed ATAC-seq peaks, and identify those that are ± 200 kb upstream and downstream of those DEGs identified in Step 1 (e.g., concordant leptin-opened peaks). Step 3: Perform motif enrichment analysis on the peaks identified in Step 2. (**B**) Pie charts showing the total number of peaks classified as fasted-opened, fasted-closed, leptin-opened or leptin-closed, as well as the subset of concordant peaks (lavender and blue bars). (**C**) TF motifs enriched in concordant fasted-opened, fasted-closed, leptin-opened and leptin-closed ATAC-seq peaks. (**D**) Heatmap showing motif scores [-log_2_(p-value)] concordant fasted-opened, leptin-closed, leptin-opened, and fasted-closed ATAC-seq peak sets. (**E**) Comparison of TF motif enrichment scores for fasted-opened peaks vs. leptin-closed peaks. (**F**) Venn diagrams showing the number of TF motifs enriched in concordant fasted-opened, leptin-closed, leptin-opened, and/or fasted-closed ATAC-seq peaks. (**G**) Summed motif enrichment scores from fasted-opened and leptin-closed TF motifs. (**H**) Comparison of TF motif enrichment scores for leptin-opened peaks vs. fasted-closed peaks. (**I**) Summed motif enrichment scores from leptin-opened and fasted-closed TF motifs.

We next performed motif enrichment analysis on the four distinct sets of concordant peaks (i.e., fasted-opened, fasted-closed, leptin-opened, leptin-closed), while employing 6 different background control peak sets. Thus, in all, 6 separate instances of AME were performed for each peak set. Motifs were considered further only if they were enriched in ≥5 out of 6 AME instances (**Figure S3A, S3E, S3F**). We observed that motifs associated with the AP-1 transcription factor family were highly enriched in concordant fasted-opened peaks, including motifs for FOSB, JUNB, ATF3, JUN, and JUND (**Figures 3C, 3D**). This finding was unsurprising given that these activity-dependent TFs have been implicated as being permissive for the physiological changes that enable neurons to increase their firing rate (Yap and Greenberg, 2018). Reassuringly, STAT3 was the most enriched TF motif in leptin-opened peaks, consistent with its established role as a key mediator of leptin-induced transcriptional regulation (**Figures 3C**).

We next set out to develop a prioritized list of TFs that might drive the expression of genes that increase AgRP neuron firing during fasting. We reasoned that such TF motifs would be enriched in concordant fasted-opened and leptin-closed peaks, but not in concordant leptin-opened or fasted-closed peaks. With this approach, we identified 101 TF motifs that were enriched in concordant fasted-opened and leptin-closed peaks (**Figure 3E**). Of these motifs, only 4 were not also enriched in concordant leptin-opened and fasted-closed peaks (**Figure 3F, 3G**). Upon ranking these motifs by their summed motif score, PRGR and ANDR were the most enriched (**Figure 3G**).

Next, we set out to develop a prioritized list of TFs that potentially drive the expression of genes that decrease AgRP neuron firing after leptin treatment. We reasoned that such associated TF motifs would be enriched in the two distinct sets of concordant leptin-opened peaks and fasted-closed peaks, but not fasted-opened or leptin-closed peaks. We identified 127 TF motifs that were enriched in both concordant leptin-opened and fasted-closed peaks (**Figure 3H**). Of these, 16 TF motifs were also not enriched in fasted-opened or leptin-closed peaks (**Figure 3F, 3I**). We then ranked these 16 TF motifs based on their summed motif scores (summed motif enrichment scores in leptin-opened and fasted-closed peaks) (**Figure 3I**). With this approach we found several motifs that corresponded to TFs that have not been linked to energy homeostasis, including interferon regulatory factor 3 (IRF3), which had the most significant concordant fasted-closed motif enrichment score, and third most significant summed motif score, behind only STAT3 and ETS2 (**Figure 3I**).

### Leptin activates neuronal IRF3

*Irf3* mRNA is expressed in AgRP neurons; we saw no change in *Irf3* levels with fasting or leptin treatment (**Figure 4A**), which was unsurprising given that *Irf3* is not nutritionally regulated in other cell types (Eguchi et al., 2011; Kumari et al., 2016). We also noted the existence of an OCR near the *Irf3* TSS (**Figure 4A**). Additionally, we detected IRF3 protein expression within the ARC of adult male mice using immunofluorescence (**Figure 4B**).

**Figure 4.**
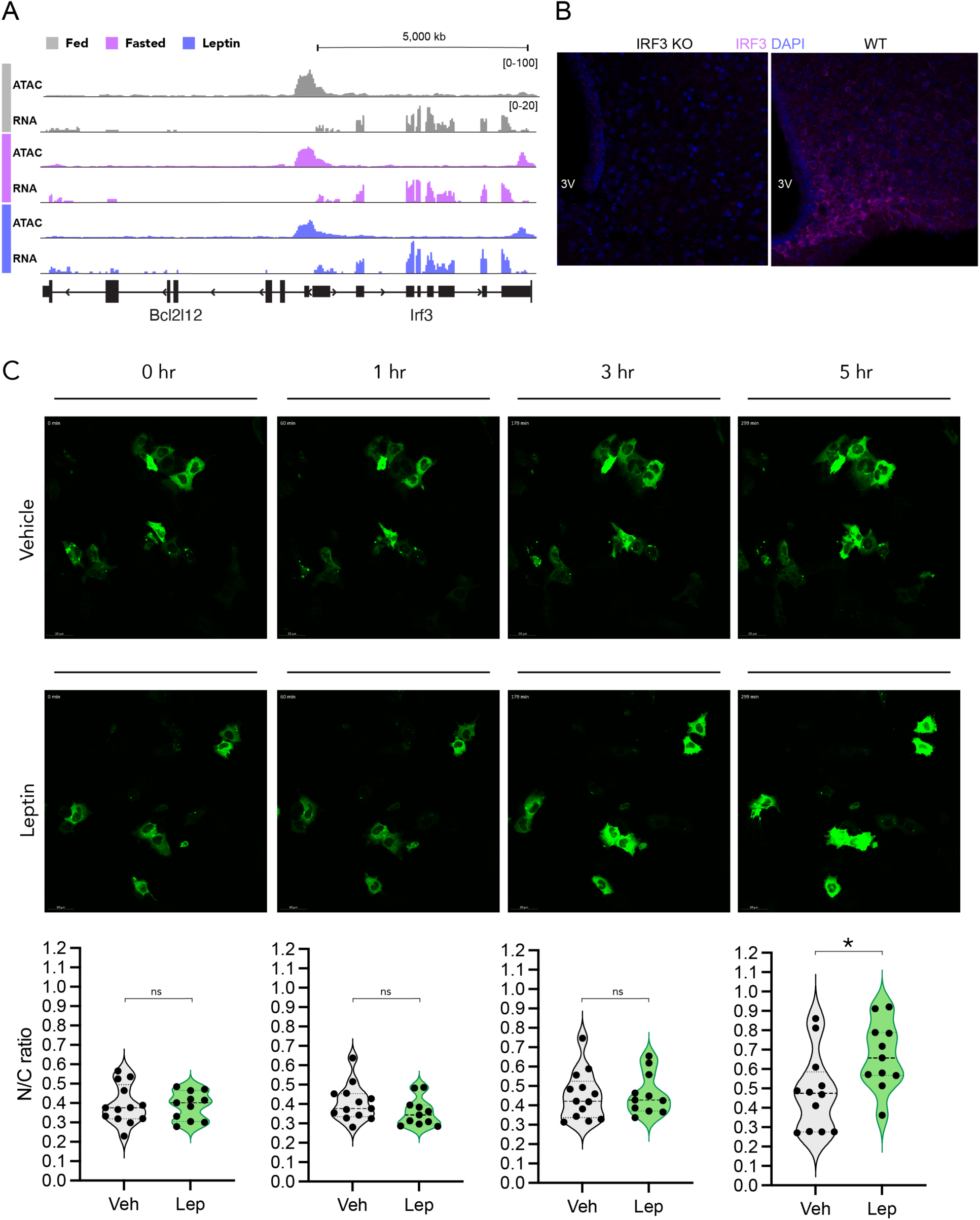
Leptin activates IRF3 in a cell autonomous manner. **(A)** ATAC-seq and RNA-seq reads at the *Irf3* locus under fed, fasted, and leptin-treated conditions. **(B)** Immunofluorescence for IRF3 (pink), with DAPI (blue), in the mouse ARC in the fed state of a WT mouse (right) and global IRF3 knockout mouse (left). **(C)** Representative images showing the dynamic imaging of GFP-tagged IRF3 in response to vehicle or leptin stimulation in GT1-7 cells expressing both IRF3-GFP and leptin receptor at 0, 1, 3, and 5 hours after stimulation. Quantification of nuclear/cytoplasmic (N/C) GFP signal is shown at each time point. Each dot represents a single tracked cell. Results are mean and analyzed by student t-test. *p < 0.05.

In immune cells, adipocytes, and other cells, IRF3 is a key component of the antiviral innate immune system and can be activated by three established pathways: (1) dsRNA → Tlr3/4; (2) dsRNA → MDA5/Rig-1; or (3) dsDNA → cGAS. Upon activation by upstream kinases, IRF3 translocates from the cytosol into the nucleus to activate gene expression (Fitzgerald et al., 2003). Interestingly, in contrast to other toll-like receptors, *Tlr3* was appreciably detected in AgRP neurons, suggesting a potential role for Tlr3 → Irf3 activation in AgRP neurons (**Figure S4A**). Moreover, in addition to Tlr3 and its signaling components, MDA5/Rig1 and their signaling components were also expressed, all of which suggest a potential route and role for Irf3 activation in AgRP neurons (**Figures S4B, S4C**).

To determine if IRF3 is activated by leptin, we employed an *in vitro* model. GT1-7 cells are an immortalized mouse gonadotrophin-releasing hormone (GnRH)-expressing hypothalamic neuronal cell line that are rendered leptin responsive when transfected with the leptin receptor (Huang et al., 2012) (**Figure S4D**). GT1-7 cells were co-transfected with plasmids expressing the long-form of the mouse leptin receptor, and IRF3-GFP, respectively. These cells were then treated with either vehicle or 100nM leptin prior to being subjected to live-cell imaging (**Figure S4E**). With this approach we observed a pronounced induction in nuclear translocation of IRF3 upon treatment with leptin at the 5-hour time point (**Figure 4C**). These results are consistent with a model whereby leptin induces the activation of neuronal IRF3 in a cell autonomous manner.

### IRF3 mediates leptin-induced satiety in AgRP neurons

We have previously observed that whole body IRF3 knockout mice are hyperphagic on a high-fat diet compared to control mice, a finding that was at the time interpreted as a compensatory mechanism to offset the increased energy expenditure that these mice exhibit (Kumari et al., 2016). Here we asked whether IRF3 mediates the hunger-suppressive effects of leptin directly. We generated male AgRP-ires-Cre::*Irf3^fl/fl^* (AgI3KO) mice as a means of deleting IRF3 selectively within AgRP neurons. No change in chow- or high fat diet (HFD)-fed body weight or cumulative food intake was apparent during basal conditions in AgI3KO mice (**Figures S5A-D**). We next implemented an established fasting-refeeding paradigm to assess leptin-induced satiety (**Figure 5A**). In this paradigm, leptin-treated AgI3KO mice exhibited greater 24-hour cumulative food intake upon refeeding than control mice, indicating blunted sensitivity to leptin (**Figure 5C**). The reduced responsivity to leptin exhibited by AgI3KO mice is unlikely to be due to general hyperphagia, or greater sensitivity to starvation, *per se*, as no difference was observed between vehicle-treated fasted-refed AgI3KO mice and control mice (**Figure 5B**).

**Figure 5.**
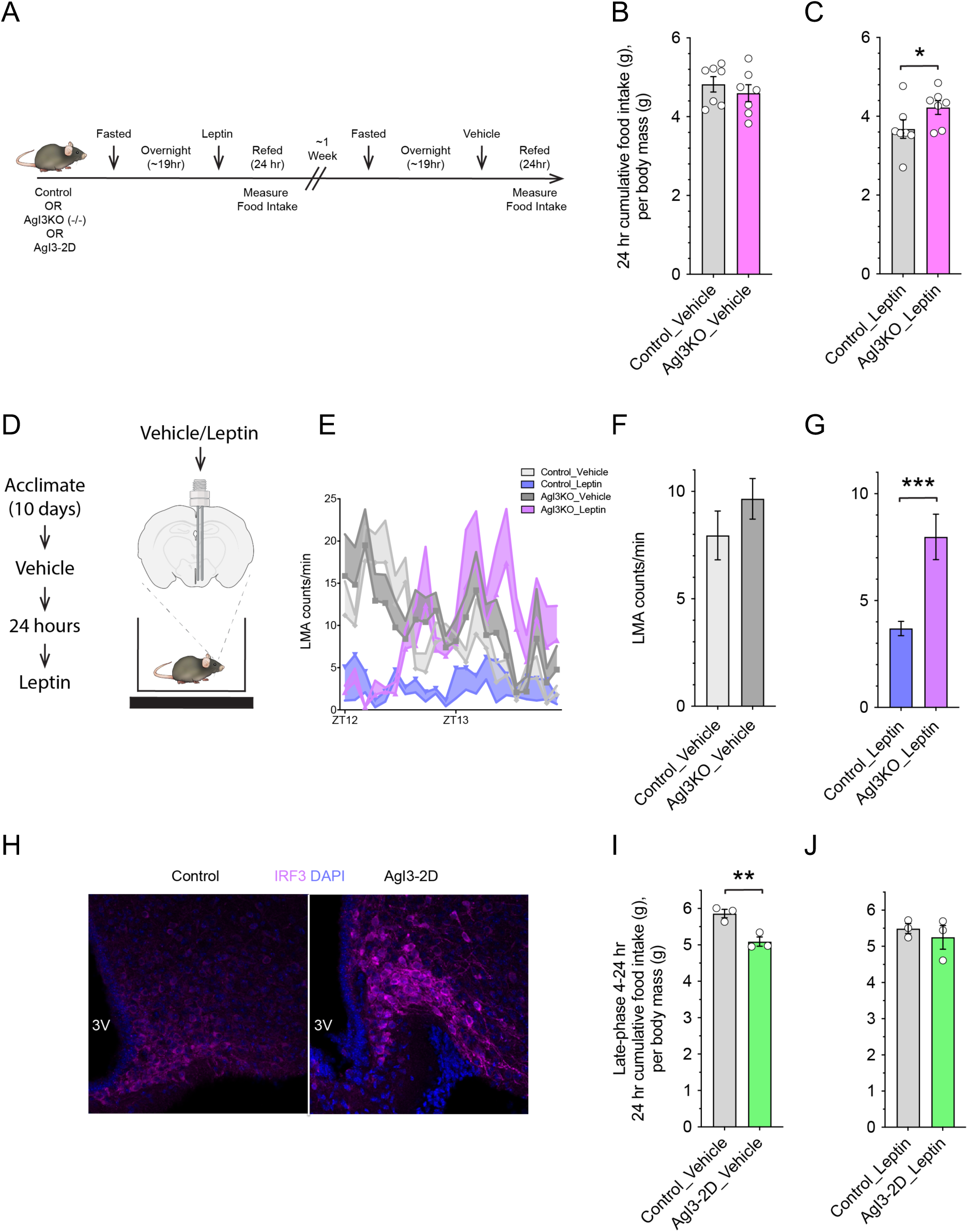
IRF3 in AgRP neurons mediates leptin-induced behavioral alterations. **(A)** Schematic of fasting-refeeding experimental paradigm. Control and AgI3KO mice were fasted overnight, followed by refeeding in the presence of leptin; food intake was measured over a 24-hour period. After a week, the same mice were subjected to the same paradigm, except vehicle was administered instead of leptin. **(B)** 24-hour cumulative body-weight adjusted food intake measured in fasted-refed, vehicle-treated, control and AgI3KO mice. **(C)** 24-hour cumulative body-weight adjusted food intake measured in fasted-refed, leptin-treated, control and AgI3KO mice. **(D)** Schematic showing the time-course and experimental paradigm of vehicle- and leptin-treatment telemetry experiments. **(E)** Locomotor activity (LMA) measurements during a 2-hour period following vehicle or leptin injected control or AgI3KO mice. (left), with quantification (right). **(F)** 2-hour LMA levels comparing vehicle-treated, control and AgI3KO mice. **(G)** 2-hour LMA levels comparing leptin-treated, control and AgI3KO mice. **(H)** Immunofluorescence of IRF3 (pink) and DAPI (blue) in the ARC for both control (left) or AgI3-2D (right) mice. (**I**) 4-24-hour body weight adjusted cumulative food intake measured by fasted-refed, vehicle-treated, control and AgI3KO mice. (**J**) 4-24-hour body weight adjusted cumulative food intake measured by fasted-refed, leptin-treated, control and AgI3KO mice. Each circle denotes a single mouse. Results are mean ± SEM of each condition and analyzed by student t-test. *p < 0.05, **p < 0.01, ***p < 0.001.

IRF3 is activated by viral RNA binding to Toll-like receptor 3 (TLR3), which is enriched in AgRP neurons relative to other TLRs (**Figure S4A**). The synthetic (TLR3) ligand poly:IC, a known activator of IRF3, induces reduced locomotion, anorexia, and early phase hyperthermia followed by a late phase (8-16 hours) hypothermia (Cunningham et al., 2007; Zhu et al., 2016). Interestingly, leptin has been shown to produce not only anorexia, but also an alteration in core body temperature that resembles a sickness response (Luheshi et al., 1999). Thus, given the high probability that AgRP neurons are a component of the neural ensemble involved in the sickness response, we asked whether IRF3^AgRP^ may mediate behavioral and physiologic responses to leptin that are commonly observed during a sickness response (i.e., reduced locomotion and derangements in thermoregulation). To this end, telemetry probes were implanted in AgI3KO and control mice to measure their locomotor activity (LMA) in two-dimensional space, as well as their core body temperature, and leptin was infused directly into the ARC (**Figure 5D**). Under vehicle-treated conditions, we noted no difference in LMA between AgI3KO and control mice (**Figure 5E, 5F**). As predicted, leptin treatment reduced LMA in control mice, but failed to do so in AgI3KO mice (**Figures 5E, 5G**). Additionally, leptin induced a modest yet significant late-phase (8-16 hours post-injection) relative hypothermia compared to vehicle (**Figures S5E, S5G**), an effect that was significantly blunted in leptin-treated AgI3KO animals (**Figures S5F, S5G**).

To assess whether IRF3 activation in AgRP neurons is sufficient to suppress food intake, we utilized a gain-of-function mouse model in which Ser388/Ser390 of IRF3 are mutated to phospho-mimetic Asp residues (S→D), creating a constitutively active allele that is expressed in a Cre-dependent manner (the allele and mouse are hereafter referred to as IRF3-2D) (**Figure S5H**). These IRF3-2D mice were crossed to AgRP-IRES-Cre mice, thereby generating AgIRF3-2D mice which expressed the constitutively active version of IRF3 in their AgRP neurons (**Figure S5H**). AgI3-2D mice exhibited an accumulation of nuclear IRF3 selectively within the ARC, a finding that is attributed to IRF3-2D being selectively enriched within AgRP neurons (**Figure 5H**). As predicted, fasted, vehicle-treated, AgI3-2D mice have suppressed food intake following a period of fasting, thus mimicking that of control mice treated with leptin (**Figures 5I, 5J**).

### Identifying transcriptional targets of IRF3 in AgRP neurons

In order to identify target genes of IRF3 in AgRP neurons that might mediate the effects of leptin on satiety, we set out to identify sites where IRF3 binds DNA, using Cleavage Under Targets & Release Using Nuclease (CUT&RUN). CUT&RUN requires less input material compared to traditional TF ChIP-seq (Liu et al., 2018; Skene et al., 2018) (**Figure 6A**). We first confirmed that we could perform CUT&RUN in neurons using sorted cerebellar nuclei to determine the genomic targets of the abundant and ubiquitously-expressed TF CCCTC-binding factor (CTCF). To this end, we conducted a titration CTCF CUT&RUN experiment using a range of 1,000-200,000 isolated cerebellar nuclei. Visual inspection of CUT&RUN data showed that all tested input levels generated high-quality profiles of CTCF peaks across the genome (**Figure S6A**). To systematically examine CTCF CUT&RUN peak signals in these titrations, we analyzed the correlation between the highest input with 200,000 nuclei and each of the lower inputs. All comparisons displayed a strong positive correlation, although correlations were weaker with lower input (**Figure S6B**). Visualization of peaks revealed highly consistent CTCF CUT&RUN peak profiles across all input depths (**Figure S6C**). Strong overlaps between peaks were observed across input levels (**Figure S6D**), with 89%-95% of peaks called in the 200,000 nuclei sample overlapping with those in the 10,000 and 50,000 nuclei samples, whereas only 47% overlapped with those in the 1000 nuclei sample. Moreover, *de novo* motif discovery confirmed that the CTCF motif was robustly enriched for each of the titration input amounts (**Figure S6A**). Next, we sought to determine whether we could perform *in vivo* CTCF CUT&RUN using 3000 AgRP neuronal nuclei, the average number we could obtain from a single NuTRAP^AgRP^ mouse. With this experiment, we observed CTCF peaks that exhibited a high coverage level and CTCF CUT&RUN peaks within genomic regions that were concordant with and distinct to those observed in the cerebellar samples (**Figure S6A**).

**Figure 6.**
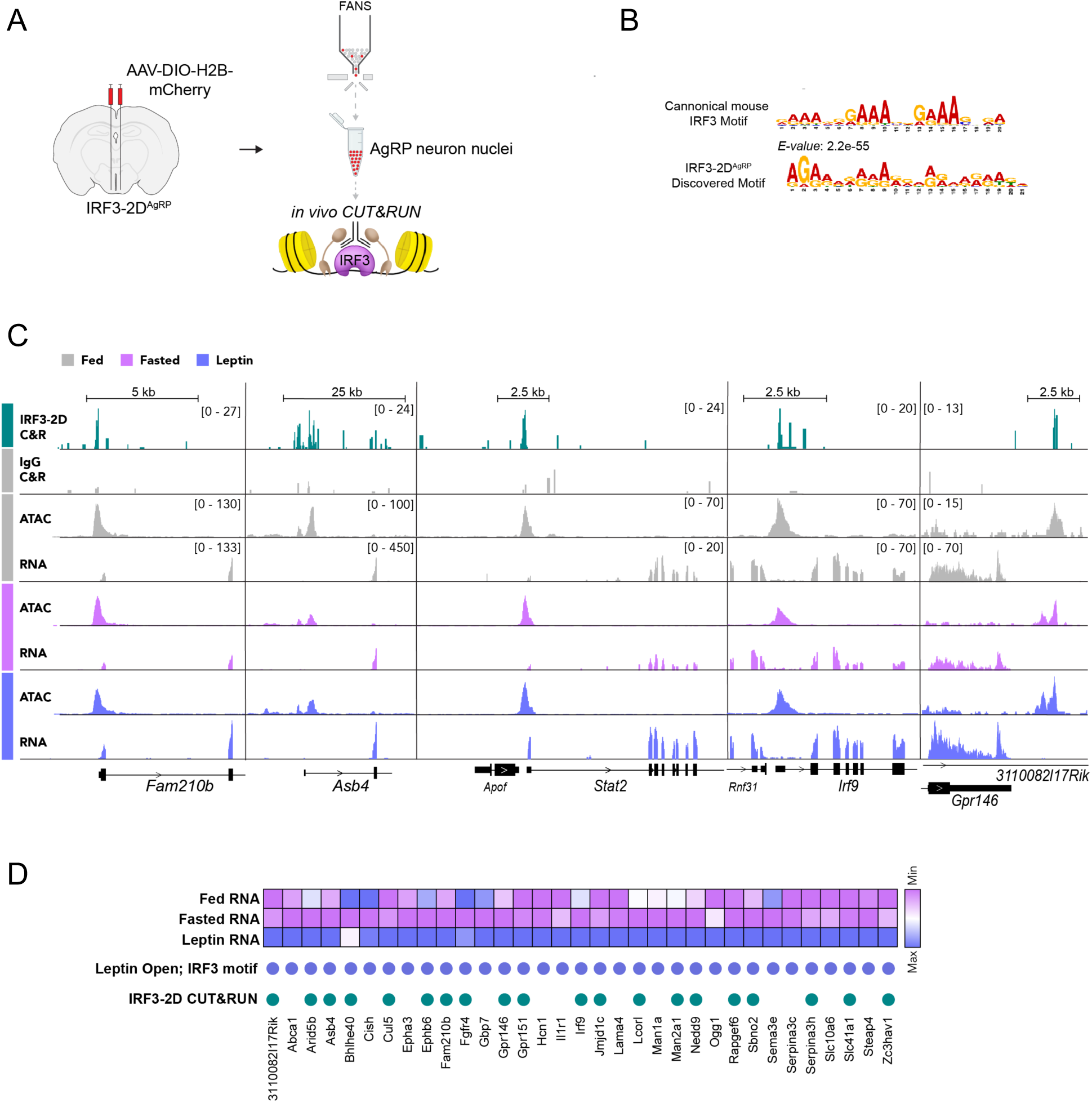
Identification of putative leptin-sensitive IRF3 transcriptional targets in AgRP neurons. **(A)** Schematic showing the *in vivo* IRF3^AgRP^ CUT&RUN approach. **(B)** Evidence of IRF3-2D^AgRP^ CUT&RUN *de novo* motif discovery analysis yielding the IRF3 motif. **(C)** Genome browser views (IGV) of IRF3 CUT&RUN peaks overlapping with ATAC-seq peaks that contain the IRF3 motif, and leptin-induced RNA-seq tracks (showcased genes from left to right: *Fam210b*, *Asb4*, *Stat2*, *Irf9*, *Gpr146*). **(D)** Heatmap showing the expression profiles of selected leptin-induced genes, with the indication of associated concordant and/or stable leptin-opened ATAC-seq peaks that contain the IRF3 motif (blue circles), and overlapping IRF3 CUT&RUN peaks (green circles).

Having validated TF CUT&RUN for low-input AgRP neuronal samples, we collected AgRP neuronal nuclei from AgI3-2D mice in the fed state and subjected them to IRF3 CUT&RUN (**Figure 6A**). In all, we detected 8,195 IRF3 CUT&RUN peaks; *de novo* motif discovery using the top 500 peaks revealed that the IRF3 motif was highly enriched (**Figure 6B**). Moreover, peaks were observed within the promoter region of suspected IRF3^AgRP^ targets (e.g., *Stat2*, *Fam210b, Irf9*) (**Figures 6C, S6E**). Of most interest, we identified 148 genes that were transcriptionally up-regulated by leptin, and contained a proximal ATAC-seq peak with an IRF3 motif that overlaps an IRF3-2D CUT&RUN peak (**Figure 6D**). Such targets include *Fam210b*, *Asb4*, *Stat2*, *Irf9*, and *Gpr146* (**Figures 6C, 6D, S6E**). Interestingly, many canonical inflammatory genes (e.g., *Isg15*, *Rsad2*) which have been linked to IRF3 activation in macrophages and other cell types were very lowly, if at all, expressed, in AgRP neurons, either in the presence or absence of leptin. Furthermore, IRF3 did not appear to bind within the vicinity of these genes (**Figures S6E, S6F**). Using ATAC-seq and IRF3 CUT&RUN data from hepatocytes as a comparator, we note that genes that are IRF3 sensitive in hepatocytes and AgRP neurons (e.g., *Stat2* and *Fam210b*) tend to display direct IRF3 binding in open chromatin regions found in both cell types (**Figure S6E**). In contrast, genes activated by IRF3 in hepatocytes but not in AgRP neurons (e.g., *Rsad2*, *Isg15*, *Ccl5*, and *Oasl2*) are associated with OCRs and IRF3 peaks in hepatocytes only (**Figure S6F**). This suggests that the underlying chromatin landscape in a given cell type determines whether IRF3 can bind and activate a particular target gene.

## DISCUSSION

While the ability of leptin to evoke prolonged changes in food consumption has largely been attributed to its engagement of the transcriptional effector STAT3, we lack a complete roster of leptin-sensitive TFs in any cell type (Beutler et al., 2017). Converging findings indicate a role for leptin-sensitive, STAT3-independent transcriptional effectors. For instance, the dose of leptin required to suppress food intake in rats is much higher than the dose required to produce maximal STAT3 activation (i.e., phospho-STAT3) within the ARC, suggesting that leptin-mediated activation of pSTAT3 is not sufficient to yield a full satiety-evoking effect (Harris, 2021). Moreover, mice lacking STAT3, or the tyrosine residue of LepR that recruits STAT3 (Tyr_1138_), are less obese than mice lacking either leptin (*ob/ob*) or the leptin receptor (*db/db*), and leptin signaling remains partially intact in these mice (Barnes et al., 2020). These results suggest the existence of a STAT3-independent arm of LepR signaling. Interestingly, LepRb sequences between residues 921 and 960 have been implicated as indispensable for the STAT3-independent component of leptin’s metabolic actions (Barnes et al., 2020). Our data indicates a role for IRF3 as a transcriptional mediator of leptin action in AgRP neurons.

Our simultaneous examination of transcriptomic and chromatin accessibility landscapes during opposing states of hunger and leptin-induced hunger-suppression enabled us to generate a list of candidate TFs that potentially regulate hunger in a leptin-dependent manner. The observation that the IRF3 motif was enriched in concordant leptin-opened peaks suggested that leptin-driven activation of IRF3 may be permissive for the expression of “hunger-suppressing” transcriptional programs in AgRP neurons. Consistent with this model, loss-of-function AgI3KO mice exhibited reduced leptin-induced suppression of hunger during a fasting-refeeding paradigm. Alternatively, or in addition, the finding that the IRF3 motif was enriched in concordant fasted-closed peaks suggested that the loss of IRF3 activity might be permissive for expression of “pro-hunger” transcriptional programs in AgRP neurons in the fasted state. Given the decline in serum leptin levels and leptin signaling during the fasted state, it stands to reason that a decline in IRF3 activity during fasting might be necessary for the establishment of a “pro-hunger” transcriptional state in AgRP neurons. Consistent with this model, gain-of-function AgI3-2D mice exhibited reduced hunger when tested in the fasting-refeeding paradigm. Given these findings, we propose that IRF3 represents a leptin-sensitive TF that bidirectionally regulates hunger, with the loss of leptin-mediated IRF3 signaling in the fasted state representing a critical “pro-hunger” event, and the resurgence in leptin-mediated IRF3 signaling in the leptin-treated state representing a key “hunger-suppressing” event.

IRF3 has been chiefly studied as a core component of the innate immune antiviral response pathway, with viral pathogen-associated molecular patterns (PAMPs) binding to TLR3/4 and activating the cGAS-STING/TBK1 pathway, resulting in phosphorylation and nuclear translocation of IRF3 (Antonczyk et al., 2019; Shu et al., 2014). The signal transduction mechanisms linking leptin receptor to IRF3 remain to be determined, but we note that several of the known upstream activators of IRF3 (e.g., cGAS, STING and IKKe) are not expressed in AgRP neurons, according to our RNA-seq results. It is interesting to speculate that PI3K-Akt, a known but poorly understood arm of the leptin signaling pathway, may be involved. This hypothesis is supported by data showing that macrophage IRF3 is phosphorylated and activated by Akt, and this activation is blocked by the PI3K inhibitor wortmannin (Joung et al., 2011; Yeon et al., 2015). Future studies will address a potential leptin → PI3K/Akt → IRF3 pathway and its contribution to leptin-induced satiety.

Mice with IRF3 ablation in AgRP neurons exhibited blunting of leptin sensitivity, yet did not exhibit either a basal or high-fat diet-induced increase in body weight. This finding is not unexpected. First, Cre-loxP deletion of *Lepr* from AgRP neurons results in only a mild increase in body mass and composition that was primarily driven by decreased energy expenditure, as no change in basal food intake was detected (Wall et al., 2008). More recently, CRISPR–Cas9-mediated genetic ablation of *Lepr* in AgRP neurons of adult mice was shown to cause an increased body weight phenotype that recapitulated that observed with whole-body *Lepr* null mice, suggesting that deletion of the leptin receptor early in development results in yet-to-be elucidated compensation that preserves the function of the core homeostatic regulatory machinery (Xu et al., 2018). A similar developmental compensation may contribute to the lack of a body weight phenotype in our AgI3KO mice. There is known functional redundancy between IRF3 and IRF7, and IRF7 or another TF (e.g., STAT3, STAT1), may be able to compensate for the absence of IRF3 (Antonczyk et al., 2019). It should be mentioned that there is precedent for Cre-loxp deletion of a suspected “pro-satiety” transcription factor abrogating leptin-induced satiety in the absence of a body weight phenotype, as was observed with the deletion of ATF3 from leptin receptor-expressing cells (Allison et al., 2018). Future studies should assess the impact of loss of IRF3 in adult AgRP neurons using a time-restricted approach. Moreover, our study was designed to identify transcriptional regulators that influence AgRP neuronal biology after an acute administration of leptin. Experiments using lean and obese mice will identify alternative candidate TFs that regulate programs underlying obesity.

IRF3 is expressed in most mammalian cells, and some of its transcriptional targets have been identified in macrophages and adipocytes (Kumari et al., 2016; Ourthiague et al., 2015; Yan et al., 2021). In immune cells, IRF3 is known to drive the expression of interferon beta (*Ifnb1*) along with a whole host of interferon stimulated genes (ISGs), cytokines, and other mediators of the innate antiviral response pathway (Antonczyk et al., 2019). Interestingly, our study revealed that many canonical IRF3 target genes were not detected in AgRP neurons during either of the two states in which we predict IRF3 is active (e.g., fed and leptin-treated), including *Ifnb1*, *Isg15*, *Isg54*, *Isg56*, *Ccl4*, *Ccl5*, *Cxcl10*, among others. Thus, IRF3 in AgRP neurons has transcriptional targets that are distinct compared to those found in other cell types. Our ATAC-seq data suggests that this may be due to an underlying chromatin landscape in AgRP neurons that restricts access of IRF3 to classical target genes. It is also possible that specific interacting proteins direct IRF3 to distinct loci in hepatocytes and immune cells vs. neurons. The IRF3 cistrome has not been determined in any other CNS cell types, and so the generalizability of this phenomenon is unclear.

It is interesting to speculate why a pro-inflammatory transcription factor might be co-opted for use in energy homeostasis. During illness, such as after viral infection, appetite is generally suppressed, an effect known as the sickness response. Polyinosinic:polycytidylic acid (Poly:IC), a synthetic analog of double-stranded RNA and a potent activator of TLR3, has been shown to mimic a viral infection, and is able to evoke a classic sickness response, including hypophagia and decreased locomotor activity, when systemically or centrally administered to mice (Cunningham et al., 2007; Zhu et al., 2016). IRF3 is a dominant transcriptional effector of TLR3 receptor activation (Antonczyk et al., 2019), and it is highly probable that IRF3 mediates the transcriptional programs underlying the prolonged sickness behaviors caused by poly:IC treatment or viral infection. It might therefore make sense for evolution to converge on the same pathway as an effector of homeostatic appetite control outside of the context of illness, such as during fasting or feeding.

## LEAD CONTACT AND MATERIALS AVAILABILITY

Further information and requests for resources and reagents should be directed to and will be fulfilled by the lead contact, Evan D. Rosen.

## EXPERIMENTAL MODEL AND SUBJECT DETAILS

### Mouse Models

#### Generation of mice

IRF3-2D mice and IRF3 floxed mouse founder lines were generated as previously described (Yan et al., 2021). For loss-of-function studies, we crossed *Irf3^flox^* mice with AgRP-IRES-Cre mice to generate AgRP neuron-deficient IRF3 mice (AgI3KO). For gain-of-function studies, we crossed *IRF3^2D^* mice with AgRP-IRES-Cre mice to generate mice expressing constitutively active IRF3 in their AgRP neurons (AgI3-2D). We crossed transgenic Nuclear tagging and Translating Ribosome Affinity Purification (NuTRAP) mice with AgRP-IRES-Cre mice to generate the NuTRAP^AgRP^ mouse line, from which we could isolate AgRP neuron-specific mRNA and nuclei.

#### Other sources

Global IRF3 knockout (IRF3KO) mice were obtained from the RIKEN BRC Experimental Animal Division (Sato et al., 2000). C57BL/6J (WT), AgRP-IRES-Cre, POMC-IRES-Cre and B6;129S6-Gt(ROSA)26Sor^tm2(CAG-NuTRAP)Evdr^/J (NuTRAP) mice were ordered from the Jackson Laboratory.

### Animals: standard fed, fasted, and leptin-treated comparison

All animal experiments were performed with approval from the Institutional Animal Care and Use Committees of The Harvard Center for Comparative Medicine and Beth Israel Deaconess Medical Center. 6-to-11-week-old male C57BL/6J NuTRAP^AgRP^ male mice were fed a standard chow diet *ad libitum*. At least one day before the experiment, mice were singly-housed in a cage with wood chip bedding and handled by the experimenter using a cupping method shown to reduce anxiety in mice (Hurst and West, 2010). Mice were either maintained on their chow diet (fed mice), or fasted (fasted mice) overnight for 18-20 hours. At zeitgeber ZT time 2-4, mice were intraperitoneally (i.p.) injected with either vehicle (PBS) or leptin (5mg/kg) and euthanized 3 hours later. As described in Campbell et al., 2017, with minor alterations, brains were rapidly extracted, and then placed ventral surface up into a chilled stainless steel brain matrix (catalog no. SA-2165, Roboz Surgical Instrument Co., Gaithersburg, MD) embedded in ice-cold PBS. Using GFP fluorescence to demarcate the ARCs rostral and caudal boundaries, brains were blocked to obtain a single coronal section containing the entire GFP+ arcuate, ∼2 mm thick. The ARC was crudely microdissected using a surgical blade at its visually approximated dorsolateral borders and immediately snap frozen on dry ice and stored at -80°C.

### Immunohistochemistry

Immunohistochemistry was performed as described (Campbell et al., 2017) with minor alterations. Mice were terminally anesthetized with 7% chloral hydrate (350 mg/kg) diluted in isotonic saline and transcardially perfused with phosphate-buffered saline (PBS) followed by 10% PFA. Brains were removed, stored in the same fixative overnight, and then transferred into 20% sucrose at 4°C overnight and cut into 40-μm coronal sections on a freezing microtome. Brain sections were washed 3 times in PBS for 10 minutes each at room temperature (RT). Sections were then subjected to antigen retrieval first by being pretreated with 1% NaOH and 1% H_2_O_2_, in H_2_O, for 20 minutes. Sections were then treated with 0.3% glycine in H_2_O for 10 minutes, followed by incubation with 0.03% sodium dodecyl sulfate for 10 minutes. Sections were then blocked for 1 hour with 3% normal goat serum in PBS/0.25% Triton-X-100. 1:100 rabbit anti-phospho-STAT3 (Tyr705) (D3A7) XP (catalog #9145S, Cell Signaling) or rabbit anti-IRF3 (catalog #4302S, Cell Signaling) was then added and incubated overnight at 4°C. The following day, sections were washed 3 times for 10 min in PBS at RT. Next, sections were treated with Alexa Fluor 647–conjugated donkey anti-rabbit (for pSTAT3 experiments, diluted 1:100; catalog no. A-31573, Thermo Fisher Scientific) for 2 h in the dark at RT. Sections were washed three times in PBS, mounted onto gelatin-coated slides (Southern Biotech), cover slipped with Vectashield Anti-fade Mounting Medium with DAPI (Vector Labs, Burlingame, CA) and sealed with nail polish. Fluorescence images were captured with an Olympus VS120 slide scanner microscope and with a confocal microscope (Zeiss LSM510 Upright Confocal System).

### Real-Time PCR Analysis

Cells or tissues were collected in Trizol reagent (Thermo Fisher), and tissues were mechanically homogenized using the TissueLyser II bead-mill (Qiagen). Total RNA was isolated using E.Z.N.A. Total RNA Kit II (Omega Bio-Tek) based on the manufacturer’s protocol. Up to 1 μg RNA was reverse-transcribed using the High-Capacity cDNA Reverse Transcription Kit (Thermo Fisher Scientific). Using 0.5 ng cDNA in a commercial SYBR Green PCR Master Mix (Thermo Fisher Scientific) and specific gene primers (see Supplementary Table 1), qRT-PCR was performed on the QuantStudio 6 Flex Real-Time PCR System (Thermo Fisher Scientific) and normalized to the housekeeping gene *Hprt* for murine studies. Analysis of qPCR data was conducted via the 2^-ΔΔCT^ method (Livak and Schmittgen, 2001).

### Translating Ribosome Affinity Purification (TRAP) Immunoprecipitation (IP)

TRAP IP was performed as previously described (Roh et al., 2017). Briefly, for each sample 4 frozen ARC tissue samples from NuTRAP^AgRP^ mice between 6 and 11 weeks of age were pooled and lysed with Dounce homogenizers in low-salt IP buffer (50mM Tris [pH7.5], 12mM MgCl2, 100mM KCl, 1% NP-40; 100 µg/ml cycloheximide, 2mM DTT, 0.2units/ml RNasin, 1x Roche Complete EDTA-free protease inhibitor). After centrifugation, the supernatant was incubated with GFP ab290 antibody (Abcam) on a rotor, then incubated again with Dynabeads Protein G (Thermo Fisher). An internal control was also collected from the lipid-free supernatant prior to antibody incubation. RNA was isolated with the RNeasy Micro Kit (Qiagen) according to manufacturer’s instructions, and then reverse-transcribed and analyzed by qRT-PCR as described above. TRAP-isolated RNA (<100ng) was treated with the Ribo-Zero rRNA removal kit (Epicentre) to deplete ribosomal RNA and converted into double stranded cDNA using NEBNext mRNA Second Strand Synthesis Module (E6111L). cDNA was subsequently tagmented and amplified for 12 cycles by using Nextera XT DNA Library Preparation Kit (Illumina FC-131). Sequencing libraries were analyzed by Qubit and Agilent Bioanalyzer, pooled at a final concentration of 12pM, and sequenced on a NextSeq500.

### RNA-seq Analysis

RNA-seq data was aligned using hisat2 (Kim et al., 2015). Reads were assigned to transcripts using feature Counts and a GRCm38 genome modified to minimize overlapping transcripts (Liao et al., 2014). Differential expression analysis of the data was performed using EdgeR (Robinson et al., 2010). Significantly different genes were required to have an average expression, across group, of > 1 cpm, a fold change (FC) > 0.5, an adjusted p-value, false discovery rate (fdr), value of < 0.05. Gene set enrichment analysis (GSEA) gene ontology (GO) for biological process was carried out with WebGestalt (WEB-based GEne SeT AnaLysis Toolkit) (Wang et al., 2017). Genes with cpm < 1 were omitted from the GSEA analysis.

### Isolation of AgRP neuronal nuclei from NuTRAP Mice

AgRP neuronal nuclei were isolated as previously described (Roh et al., 2017), with minor alterations. Briefly, dissected ARCs from 6-11-week-old mice were snap frozen and stored at -80C. Isolated ARCs were dounce homogenized in nuclear preparation buffer (NPB; 10 mM HEPES [pH 7.5], 1.5mM MgCl_2_, 10 mM KCl, 250 mM sucrose, 0.1% NP-40, and 0.2 mM DTT), and homogenates were filtered through a 100 μM strainer and centrifuged to pellet the nuclei. Nuclei were washed with NPB, re-suspended in nuclear sorting buffer (10 mM Tris [pH 7.5], 40 mM NaCl, 90 mM KCl, 2 mM EDTA, 0.5 mM EGTA, 0.1% NP-40, 0.2 mM DTT), and filtered again through a 40 μM strainer. Isolated nuclei were sorted by flow cytometry based on AgRP neuron-specific GFP expression. With this approach we routinely obtain ∼3000 AgRP neuronal nuclei per mouse.

### Nuclei processing and library preparation for ATAC-seq

NuTRAP^AgRP^ ARC nuclei from 4 pooled mice per condition were isolated, subjected to FANS, and collected into 500-750uL PBS (0.1% NP40) in 1.5 mL microcentrifuge tubes and stored on ice. Tubes were spun at 1000rpm for 10min, at 4°C. Supernatant was removed via gentle decantation. Nuclei pellets were subjected to tagmentation after resuspension in 50ul of transposase reaction mix: 25 ul TD buffer, 2.5 ul TN5 transposase, 22.5 ul dH_2_0. Nuclei were incubated at 37°C with shaking at 600 rpm for 30 mins, then placed on ice. DNA was purified using a Qiagen minelute PCR purification kit and eluted in 10 ul of Qiagen Elution Buffer. Library construction was conducted using an initial PCR amplification using 10 ul of transposed DNA, 11ul H_2_O, 2ul of 25uM Primer 1, 2ul of Primer 2, and 25 ul of NEBNext HF 2X PCR mix. The following initial amplification cycle was used: 1 cycle or 72°C (5min), 98°C (30s), 5 cycles of 98°C (10s), 63C (30s), 72°C (1 min), hold at 4°C. A side PCR reaction was conducted to determine the optimal number of amplification cycles used during the library construction, as previously described(Buenrostro et al., 2015), using 5 ul PCR amplified DNA, 4.41 ul H_2_O, 0.25ul Primer 1 Adaptor, 0.25ul Primer 2 Adaptor 2 (barcoded), 0.09 ul 100x SYBR Green (Diluted from 10,000X stock with H_2_O), 5 ul NEBNext HF 2X PCR mix. The following initial amplification cycle was used: 1 cycle of 98°C (30s), 20 cycles of 98°C (10s), 63°C (30s), 72°C (1 min). Amplification curves were assessed to determine the number of additional cycles needed during library construction(Buenrostro et al., 2015). The requested additional number of cycles was completed by adding the sample back to the thermocycler in its same tube and master mix. Amplified libraries were purified with a Qiagen Minlute PCR purification kit, eluted in 20ul, and the volume was brought up to 100 ul with additional EB. The eluent was subjected to a two-phase size selection using Ampure Beads, initially .55X volume of elutant (55ul) was used, followed by 1.5X volume of initial sample volume (100ul x 1.5 = 150ul) minus the existing PEG that is already in the sample (150ul – 55ul = 95 ul). Finally, the library was eluted using 50ul of EB. Libraries were sequenced on a Next-seq 500 (Illumina).

### ATAC-seq data processing

Library reads were mapped to the UCSC build mm10 assembly of the mouse genome using Bowtie2, with peak calling using MACS2. ATAC-seq peaks were visualized in the WashU Epigenome Browser. Coverage summation and additional data parsing were carried out using Bedtools. Differential binding analysis including normalization and quantification were done in EdgeR. Differential peaks were defined at log2 fold change ≥ 0.5 and a false discovery rate of < 0.05.

### Motif Enrichment Analysis and TF Prioritization

Transcriptionally concordant ATAC-seq peaks were subjected to Motif Enrichment was determined using the MEME-suite Analysis of Motif Enrichment (AME) (Version 5.1.0) sequence analysis tool (**Figure 3A**)(Bailey et al., 2015). We restricted our analysis to significant concordant peaks for each of the four significant ATAC-seq peak patterns (i.e., fasted-opened, fasted-closed, leptin-opened, leptin-closed). AME was conducted 6 distinct times using one of the 6 possible background control sets of peaks during each analysis: (1) neutral (i.e., unchanging) peaks anywhere in the genome; (2) neutral peaks near neutral genes; (3) neutral peaks near significant genes; (4) neutral peaks near any TSS; (5) all non-concordant ATAC-seq peaks for a given conditions (e.g., leptin-opened); or (6) concordant ATAC-seq peaks that are shuffled (e.g., leptin-opened peak sequences that are shuffled). Thus, in all, 6 distinct instances of AME were performed for each peak set. We next employed a rule whereby we further considered motifs only if they were significantly enriched in ≥5 out of 6 AME instances (**Figure S3A, S3E**).

To determine the representative log transformed p-values (for plotting purposes), for those “enriched” motifs (at least 5 out of 6 AME instances), the minimal (most significant) p-value out of the 6 motif instances was determined, while omitting from the analysis any non-significant p-values, for those Motifs that have a single non-significant motif but are retained. Conversely, to determine the representative p-value for those non-enriched motifs (violate the at least 5 out of 6 significant rule) we calculated the minimal (most significant) p-value, of the non-significant ones. We then log transformed the resulting p values (log_2_). For the transcriptionally concordant fasted-opened condition only, p-values for 12 of the 356 TF motifs were less than e^-300 and thus could not be log transformed (ATF3.0.A, BACH1.0.C, BACH2.0.A, BATF.0.A, CRX.0.A, FOS.0.A, FOSB.0.A, FOSL1.0.A, FOSL2.0.A, JUN.0.A, JUNB.0.A, JUND.0.A). Therefore for these motifs only in fasted-opened condition, the log transformed p-value was determined by imputation. Evidently, none of these 12 TF motifs were qualified (due to enrichment in disqualifying chromatin conditions) to be plotted in figures 3G or 3I. All plots were made using Graphpad PRISM software (Version 9).

### *In vitro* leptin-induced pSTAT3 measurement experiments

Immortalized hypothalamic GT1-7 cells^36^ were plated in DMEM with high glucose (Sigma, D5796), Penicillin-Streptomycin, and 10% fetal bovine serum. Cells were transiently transfected using Lipofectamine 3000 Reagent (Thermo Fisher) based on manufacturer’s instructions. Briefly, dsDNA-lipid complexes were prepared using 0.5ug of plasmid expressing the murine long-form leptin receptor (LepR), or both LepR plasmid and the GFP-tagged IRF3 plasmid, and Lipofectamine 3000 with P1000 reagent in Opti-MEM Medium (Thermo Fisher), and then added dropwise onto cells in medium. Cells were washed with PBS, treated with serum-free DMEM, and treated with either leptin (100 nM) or vehicle (control) for 15 minutes.

### Immunoblotting

Lysis of frozen tissue was performed in 1X RIPA lysis buffer (Millipore Sigma) with the addition of protease and phosphatase inhibitors (Millipore Sigma). Tissues were mechanically homogenized using the TissueLyser II bead-mill (Qiagen). Total protein was quantified via BCA Protein Assay Kit (Thermo Scientific), protein lysate was prepared in 1X SDS in RIPA lysis buffer, and then boiled. Lysis of cells was performed as previously described (Silva et al., 2018). Briefly, media was aspirated from plated cells, and then 1X SDS in RIPA lysis buffer was added onto the cells, which were then collected, flash frozen, and boiled. The prepared protein samples were separated on Tris-HCl protein gels (Bio-Rad) and transferred to PVDF membranes (Thermo Fisher), followed by blocking in 5% blotting-grade milk (Bio-Rad). Blots were incubated in appropriate primary antibodies (1:1000 dilution unless otherwise indicated) at 4°C overnight, and then in secondary HRP-conjugated antibody (Cytiva) for one hour at RT. Blots were developed with chemiluminescent ECL reagents (Thermo Fisher) and imaged with ImageLab^TM^ Touch Software (Bio-Rad) on the ChemiDoc Touch Imaging System (Bio-Rad).

### *In vitro* IRF3 live-cell imaging and quantification

80,000 GT1-7 cells were plated in two wells of an 8-well Nunc™ Lab-Tek™ II Chambered Coverglass (Catalog # 155360) in media. The following day cells were transfected with plasmids expressing mouse LepR and IRF-GFP, as described above. The following afternoon, cells were washed with 37°C PBS and serum-free DMEM (Sigma #D1145) was added to cells prior to imaging approximately 1 hour later. Ideal cells, for which the nucleus was well demarcated, and for which the IRF3-GFP expression was apparent, were targeted and images were scanned and recorded every 10 minutes obtained using a Zeiss LSM 880 upright laser scanning confocal microscope in 3 × 3 tile-scan mode with a Plan-Apochromat 20×/0.8 M27 objective. Mean Fluorescence Intensity of each targeted cell’s cytoplasm and nuclei was determined using ImageJ software (NIH) to enable calculation of the nuclear-to-cytoplasm (N/C) GFP intensity ratio.

### Fasting-refeeding studies

6-to-11-week-old male C57BL/6J NuTRAP^AgRP^ male mice were fed standard chow diet *ad libitum*. At least one day before the experiment, mice were singly housed and handled by the experimenter using a cupping method shown to reduce anxiety(Hurst and West, 2010). Depending on the experiment, mice were fasted overnight starting at ZT 8-10. The following day, between ZT 3-5, mice were injected i.p. with leptin, and 30 minutes later a weighed food grate was given. Food was measured during a 24-hour period. For the AgI3KO fasting-refeeding experiment, a white paper mesh (Alpha pads) was added to the cage in place of bedding, to allow for the accounting of spilled food when calculating food intake. Food intake was obtained by subtracting remaining food, including any spilled food in cages, from the previously weighed food amount (Yang et al., 2014). Food intake adjusted for body weight was calculated by dividing the absolute food intake by body weight and multiplying the result by the average body weight of all mice (Bachmanov et al., 2002).

### Telemetry experiments

All surgeries were performed in sterile conditions. Mice were anesthetized with a mix of ketamine/xylazine (100 and 10 mg/kg, respectively, IP) with additional doses of 10% of the initial dose throughout surgery to eliminate the withdrawal reflex. Mice were implanted with a radiotelemetry temperature and locomotion sensor (TA-F10, DSI) in the intraperitoneal space via laparotomy(Machado et al., 2018). Meloxicam treatment, for analgesia, was administered prior to surgery, and then again 24hrs later. Mice were allowed to recover at least 10 days prior to experimentation. Following recovery, mice used for experiments showed no signs of discomfort and gained weight normally(Machado et al., 2018). Core body temperature and locomotion were recorded using the radiotelemetry DSI system. The signal was sent from the telemetry probes previously implanted to receivers and converted using the PhysioTel HD and PhysioTel (DSI) hardware which provides the mean of Tc every 5 min. All animals had at least 48 hours of acclimation in the recording chamber before baseline was recorded.

### High Fat Diet Feeding

Mice were maintained at 12h/12h light/dark cycles, 23°C room temperature and 30-70% humidity with *ad libitum* access to food and water, in individually ventilated cages. At approximately 6 weeks of age, male mice were placed on either a Standard Chow Diet (SCD; 8664 Harlan Teklad, 6.4% wt/wt fat), or a high fat diet (HFD) consisting of 20% calories from protein, 60% from fat, and 20% from carbohydrate (Research Diets, D12492) for the indicated durations. Body weight was measured weekly.

### *General In vivo* Stereotaxic Injections

Mice were anesthetized with xylazine (10mg/kg, i.p.) and ketamine (100 mg/kg, i.p.) and placed in a stereotaxic apparatus after assuring proper anesthesia effects with a toe pinch or tail pull. The fur around the skull was shaved and the skin sanitized. An incision in the skin overlying the skull was made and the skull surface exposed. A small hole was then drilled into the skull to expose the brain surface. The ARC was bilaterally localized according the coordinates listed in mouse brain in stereotaxic coordinates by Paxinos (Konsman, 2003) and the tip of the nanoject injector connected to a sterile glass pipette was be lowered into the ARC bilaterally (coordinates for ARC injections were anterior -1.37 mm, lateral ± 0.3 mm, and ventral 5.70-5.80). pAAV-hSyn-DIO-H2B-mCherry (Addgene plasmid # 50459), was then slowly injected. Afterwards, the edges of the incision were reapposed with tissue adhesive (vetbond, n-butyl cyanoacrylate). Analgesia was given in the form of Meloxicam SR (4mg/kg). Animals were allowed to recover from surgery for 2 weeks and their body weight and health conditions were closely monitored during recovery. Coordinates and injection volume used in the studies were: the ARC (anterior-posterior (AP): −1.40 mm, dorsal-ventral (DV): −5.80 mm, left-right (LR): ± 0.30 mm, 150 nl/side). Accuracy of injection was determined by the expression of mCherry driven by AgRP-IRES-Cre exclusively within the ARC, both during tissue collection and during FACS.

### Cleavage Under Targets and Release Using Nuclease (CUT&RUN)

Male AgI3-2D mice were stereotaxically injected with pAAV-hSyn-DIO-H2b-mCherry into their ARCs, and tissue isolated, as previously described. ∼4K ARC nuclei were isolated and subjected to FANS, as described above, and collected into 0.75 mL of C&R Wash Buffer (low salt) buffer (20 mM HEPES-KOH [pH 7.5], 75 mM NaCl, 0.5 mM Spermidine, 0.1% BSA and 1x protease inhibitor cocktail from Sigma). 180 μL BioMagPlus Concanavalin A beads slurry was washed twice and re-suspended in 180 μL Binding Buffer (20 mM HEPES pH 7.5, 10 mM KCl, 1 mM CaCl_2_, 1 mM MnCl_2_), and added to the nuclear suspension. The mixture was thoroughly mixed on a rocker for 10 minutes to allow binding of nuclei to the beads. The nuclei were pelleted with a magnet stand and blocked with Wash Buffer (20 mM HEPES-NaOH pH 7.5, 75 mM NaCl, 0.5 mM Spermidine, 0.1% BSA and 1x protease inhibitor cocktail from Sigma) supplemented with 2 mM EDTA. After washing once with Wash Buffer, nuclei were re-suspended in 50 μL Wash Buffer containing 2 μg of anti-IRF3 antibody (Proteintech, 11312-1-AP), and incubated overnight to allow antibody binding. The next day, nuclei were washed once on a magnet stand with Wash Buffer and incubated with 1:1000 pA-MN in 200 μL Wash Buffer. After 1 hour, nuclei were washed once to remove unbound pA-MN, re-suspended in 150 μL Wash Buffer, and then chilled on a 0°C metal block in a water-ice mixture. 3 μL of 100 mM CaCl_2_ was added to activate pA-MN, and the mixture was incubated at 0°C for 60 min. The reaction was stopped by adding 150 μL 2XSTOP buffer (200 mM NaCl, 20 mM EDTA, 4 mM EGTA, 50 μg/mL RNase A and 40 μg/mL glycogen). The protein-DNA complexes were released by centrifugation and then digested by proteinase K at 50°C overnight. DNA was extracted by ethanol precipitation, followed by Qubit fluorometer and bioanalyzer quality control. CTCF optimization experiments were performed using DAPI-treated nuclei isolated from whole mouse cerebellum, and a CTCF antibody from (Millipore, 07-729). Peri-hypothalamus IRF3-2D IgG (IgG C&R, Figures 6, S6E, S6F) sample was generated by stereotaxically injecting a IRF3^LSL-2D^ mouse with a AAV2-Cre-GFP in the peri-hypothalmus and ∼200K GFP+ nuclei were isolated by FANS and used for CUT&RUN on the same day the IRF3^Ag^ C&R sample was generated. IRF3 CUT&RUN from hepatocytes was conducted using primary hepatocytes isolated from IRF3^LSL-2D^ mice as described, and transiently transduced *ex vivo* with adenovirus expressing Cre recombinase (Ad-Cre). IRF3 and IgG CUT&RUN from immortalized AgRP neurons (IRF3 C&R AgRP*^in vitro^* and IgG C&R AgRP*^in vitro^*, Figures S6E, S6F) was conducted using those cells transiently transduced with Ad-CMV-IRF3-2D-hPGK-eGFP (Vector Builder).

### CUT&RUN Library Preparation

CUT&RUN libraries were prepared using the NEB Ultra II library preparation kit with important modifications(Liu et al., 2018). Briefly, less than 30 ng DNA was used as input. End prep was performed at 20°C for 30 minutes and then 50°C for 60 minutes, as reduced temperature prevents melting of short DNA fragments (Liu et al., 2018). 5 pmol adaptor was added and ligated to end prep products at 20°C for 15 minutes. USER^TM^ enzyme was then used to cleave the uracil in the loop. 1.7x volume of AMPure beads were used to purify the ligation product, which aids in the recovery of short fragments. To amplify the library, the ligation product was mixed with 2x Ultra II Q5 mix, universal primer and index primers. PCR was carried out as follows: 98°C 30 seconds, 12 cycles of 98°C 10 seconds and 65°C 10 seconds, and final extension 65°C 5 minutes. The PCR product was subjected to double size selection with 0.8x and then 1.2x AMPure beads. The purified library was quantified with Qubit and Tapestation. Libraries were pooled at similar molar amounts and sequenced using Nextseq500 platform. Paired-end sequencing, read length 42 bp x 2, 6 bp index was performed. The detailed protocol is given at https://dx.doi.org/10.17504/protocols.io.bagaibse

### CUT&RUN data processing

For IRF3 CUT&RUN, raw data quality check was done using fastQC then peaks were aligned to mm10 using bowtie2. Peaks were called using SEACR (Meers et al., 2019) and filtered. Motif analysis was performed using MEME-Suite by taking a radius of 100 bp around the summit. For CTCF CUT&RUN, the quality of the raw reads was checked using fastp v0.20.0, then the peaks were aligned to GRCm38 using bowtie2 v2.3.5. Peaks were called using macs2 v2.2.6 in strict (q-value 0.1) and relaxed (p-value 0.01) modes; peaks were then filtered using common peaks found in both pseudoreplicates (using p-value 0.01). For this peak list, sequence regions containing a radius of 100 base pairs around the summit was used for *de novo* motif discovery analysis using meme-chip v5.1.0. The motifs were then searched against the HOCOMOCO v11 mouse database.

## QUANTIFICATION AND STATISTICAL ANALYSIS

Sample size, mean, and significance P values are indicated in the text, figure legends, or Method Details. Error bars in the experiments represent standard deviation (STD) from either independent experiments or independent samples. Statistical analyses were performed using GraphPad Prism. Detailed information about statistical methods is specified in figure legends or Method Details.

## DATA AND CODE AVAILABILITY

All raw and processed RNA-Seq, ATAC-seq, as well as CTCF and IRF3 CUT&RUN data will be deposited in the NCBI Gene Expression Omnibus prior to publication and assigned an accession number GEO: XXXXX.

## ACKNOWLEDGEMENTS

This work was supported by an NIH 3 T32 DK 7516-32 to F.D.H., American Heart Association Postdoctoral Fellowship 18POST33990061 to F.D.H, American Diabetes Association Minority Postdoctoral Fellowship 1-19-PMF-008 to F.D.H., a Burroughs Wellcome Fund Postdoctoral Enrichment Program (PDEP) grant to F.D.H., a Nutrition and Obesity Research Center of Harvard (NORCH) URM Pilot and Feasibility Grant to F.D.H., and R01 DK085171, R01 DK102173, R01 DK102170, and R01 DK1113669 to E.D.R. We thank the BNORC Functional Genomics and Bioinformatics Core, the BIDMC Confocal Imaging Core, and the Molecular Biology Core Facilities (DFCI). We thank Steve Henikoff for providing pA-MNase. We thank Young-Bum Kim for providing GT1-7 cells and LepR plasmid. We thank Bradford Lowell (BIDMC; HMS) and Clifford Saper (BIDMC; HMS) for helpful comments regarding this manuscript.

## AUTHOR CONTRIBUTIONS

F.D.H. conceived this study and interpreted the results of all experiments. E.D.R supervised this study. F.D.H, optimized, designed, and executed TRAP-seq experiments. F.D.H and E.D.R conceived of ATAC-seq studies, which F.D.H optimized, designed, and executed with advice from E.D.R and L.T.. F.D.H designed all CUT&RUN experiments. N.L., F.D.H and S.J.P optimized initial *in vitro* IRF3 CUT&RUN. N.L. and F.D.H optimized and executed all *in vivo* CUT&RUN experiments with input from S.H.O.. C.J., R.I., and H.S., performed computational and bioinformatic analyses involving sequencing data with advice from L.T.. F.D.H conceived of motif filtering/prioritization scheme. C.J. and F.D.H curated RNA-seq, ATAC-seq, and Motif enrichment analyses. C.J. curated integrated IRF3 CUT&RUN data. F.D.H performed data visualization for all experiments with the exception of the CTCF CUT&RUN experiment. H.S. performed data curation and visualization for CTCF CUT&RUN experiments. F.D.H conceived of, designed, and executed *in vitro*, live-cell imaging, and western blot, experiments. F.D.H optimized, designed, and executed all *in vivo* immunofluorescence experiments. F.D.H, N.M., and A.U. designed and executed telemetry experiments. F.D.H wrote the manuscript with major input from E.D.R. along with added input from L.T., and all other authors.

---

**Supplementary Figure 1.**
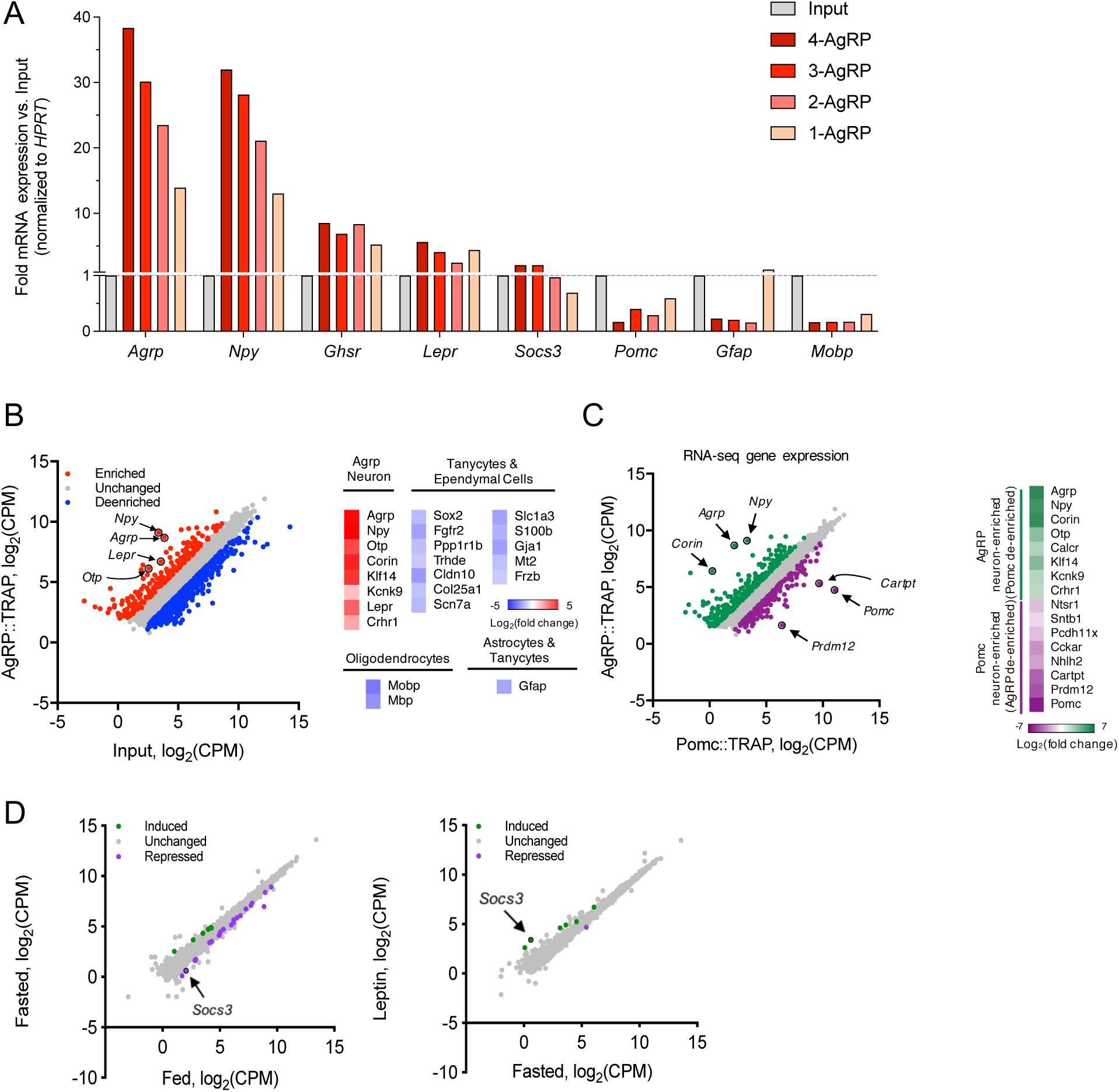
Establishment of TRAP-seq for AgRP neurons. Associated with main Figure 1. **(A)** Gene expression analysis by qPCR with TRAP-isolated RNA from a titration of 4-to-1 pooled NuTRAP^AgRP^ ARCs. Bars indicate the fold mRNA expression vs. input for each sample. **(B)** Scatter plot of RNA-seq data showing regulated genes from TRAP-enriched AgRP neurons versus input control fraction of these same samples (*n* = 3, 3) [fold change > 0.5 up (red) or down (blue) and false discovery rate (FDR) < 0.05]; CPM, counts per million. The number of genes in a given expression profile is in parentheses. **(C)** Scatter plot of RNA-seq data showing regulated genes within AgRP neurons from TRAP-enriched AgRP neurons versus Pomc neurons from TRAP-enriched Pomc neurons (*n* = 3, 2) [fold change > 0.5 up (red) or down (blue) and false discovery rate (FDR) < 0.05]; CPM, counts per million. **(D)** Left: Scatter plot of RNA-seq data showing regulated genes within Pomc neurons from fasted versus fed littermates (*n* = 2, 2) [fold change >0.5 up (red) or down (blue) and false discovery rate (FDR) < 0.05]; CPM, counts per million. Right: Scatter plot of RNA-seq data showing regulated genes within Pomc neurons from leptin-treated versus fasted littermates (*n* = 2, 2) [fold change >0.5 up (red) or down (blue) and false discovery rate (FDR) < 0.05]; CPM, counts per million.

**Supplementary Figure 2.**
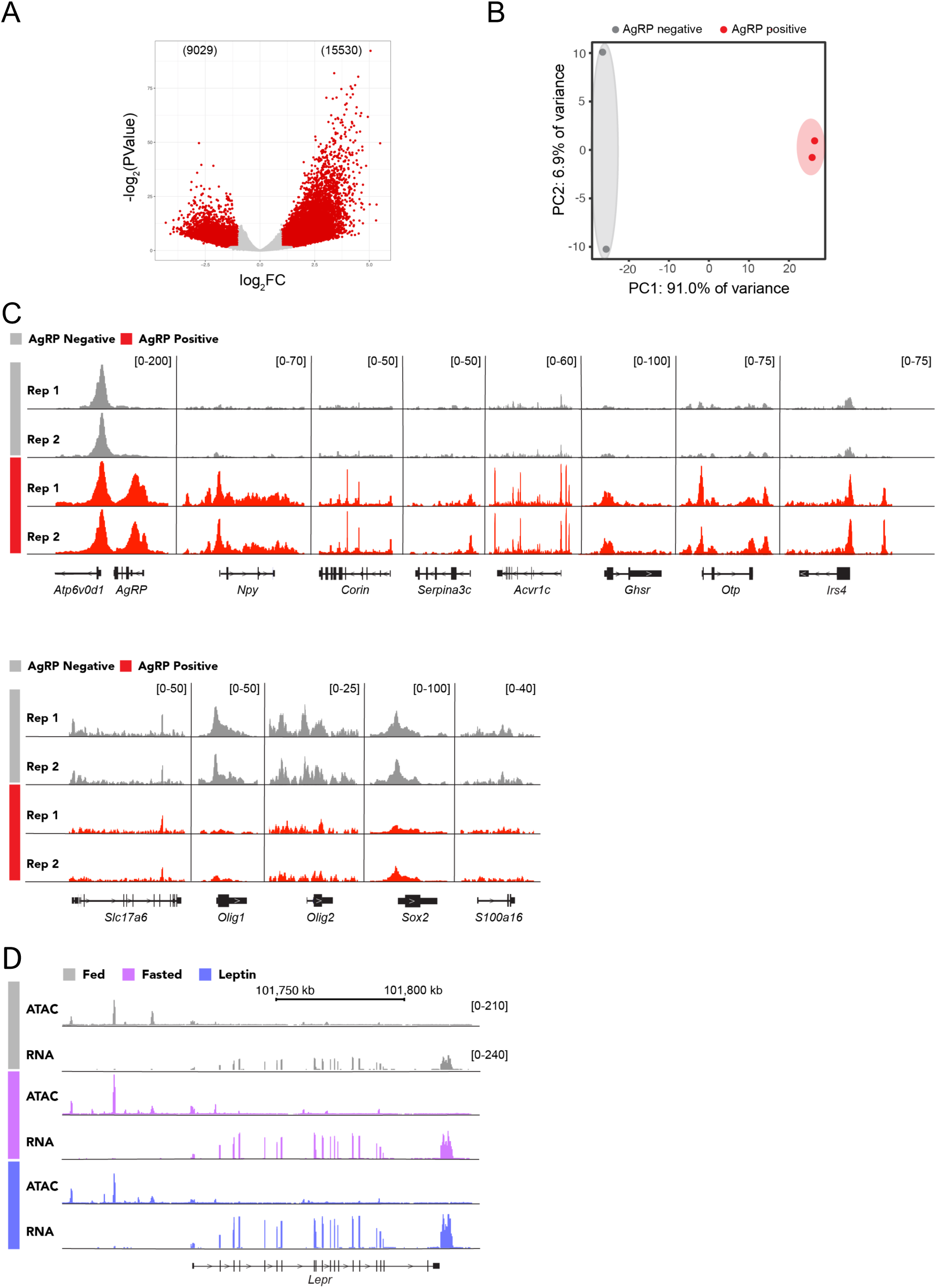
ATAC-seq profiles of AgRP neurons. Associated with main Figure 2. (**A**) Volcano plot showing differential OCRs upon comparing AgRP positive vs. AgRP negative (n = 2, 2) neurons. Red dots correspond to significantly different gained-open and gained-closed regions [fold change >0.5 up (red) or down (red) and false discovery rate (FDR) < 0.05]; CPM, counts per million. (**B**) Principal Components Analysis (PCA) plot comparing AgRP negative and AgRP positive ATAC-seq samples. (**C**) Genome browser views (IGV) of representative ATAC-seq peaks near genes enriched (top) and de-enriched (bottom) in AgRP neurons. (**D**) Genome browser views (IGV) of *Lepr*-associated ATAC-seq and RNA-seq peaks in the fed, fasted, and leptin-treated states.

**Supplementary Figure 3.**
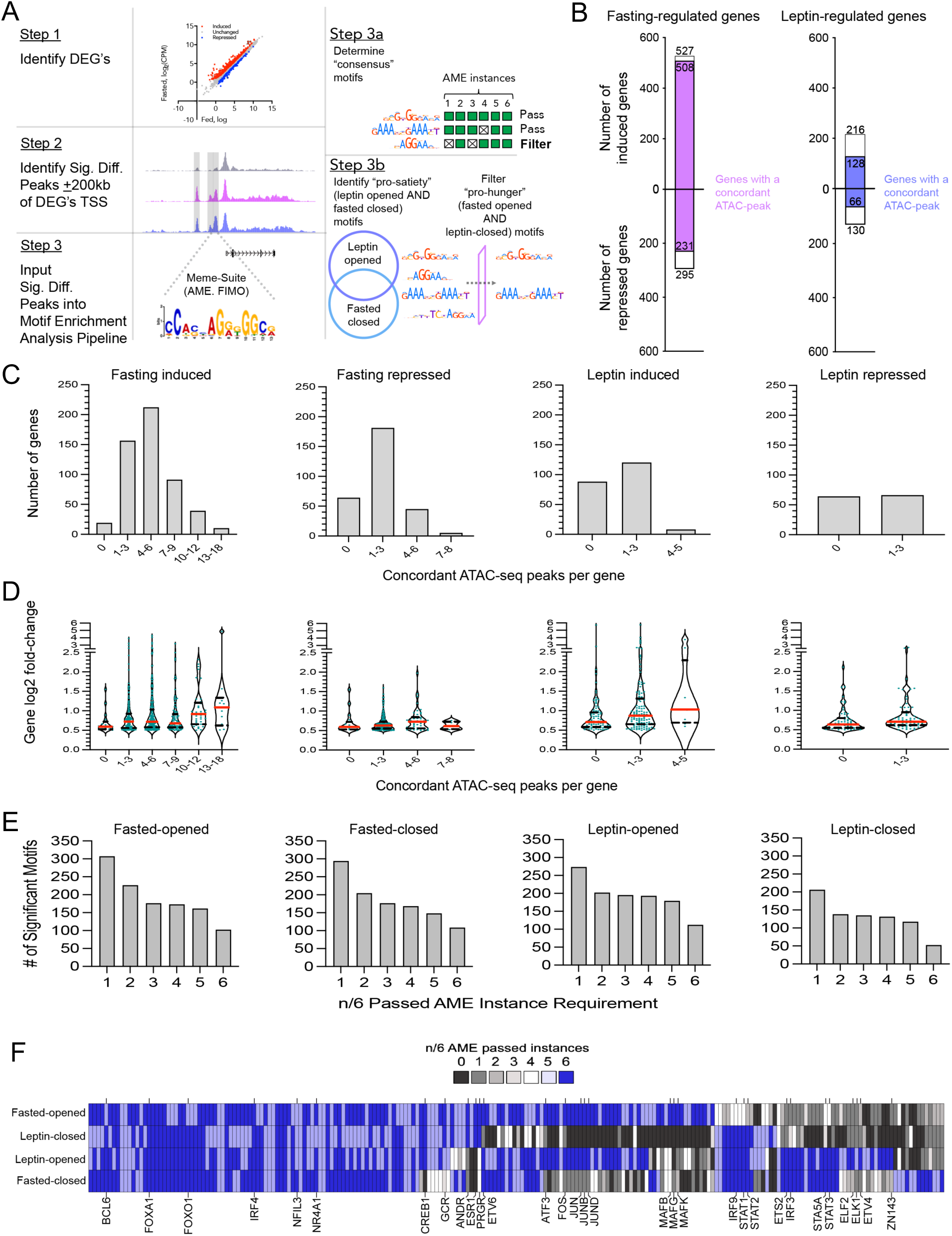
Gene expression and chromatin state metrics of AgRP neurons. Associated with main Figure 3. (**A**) Schematic showing the detailed computational heuristic employed to identify putative pro-satiety TF motifs. Step 1: Identify fasted- and leptin-induced differentially expressed genes (DEGs). Step 2: Determine significantly different fasted-opened, fasted-closed and leptin-opened ATAC-seq peaks, and identify those that are ± 200 kb upstream and downstream of those DEGs identified in Step 1 (e.g., gene-linked ATAC-seq peaks). Step 3a: Perform motif enrichment analysis on the peaks identified in Step 2. Step 3b: Determine the consensus motif identified in 5 out of 6 instances of analysis of motif enrichment (AME). Step 3c: Identify putative pro-satiety TF motifs that are dually enriched in leptin-opened and fasted-closed peaks, but not fasted-opened (putatively pro-hunger) peaks. **(B)** Left: bar graphs showing number of fasted-induced and fasted-repressed genes, with internal bar showing the number of those genes that have a concordant ATAC peak. Right: bar graphs showing number of leptin-induced and leptin-repressed genes, with internal bar showing the number of those genes that have a concordant ATAC peak. **(C)** Number of genes that possess a given number range of concordant ATAC-seq peaks per gene for fasting-induced, fasting-repressed, leptin-induced, leptin-repressed conditions. (**D**) Mean gene fold-change (red bar) for binned numbers of concordant ATAC-seq peaks per gene for fasting-induced, fasting-repressed, leptin-induced, leptin-repressed conditions. (**E**) Number of significant motifs for each of the passed AME instance requirements (1-6), for fasted-opened, fasted-closed, leptin-opened, and leptin-closed conditions. (**F**) Heatmap of the number of AME instances that passed the significance threshold (i.e., n ≥ 5 out of 6 AME instances).

**Supplementary Figure 4.**
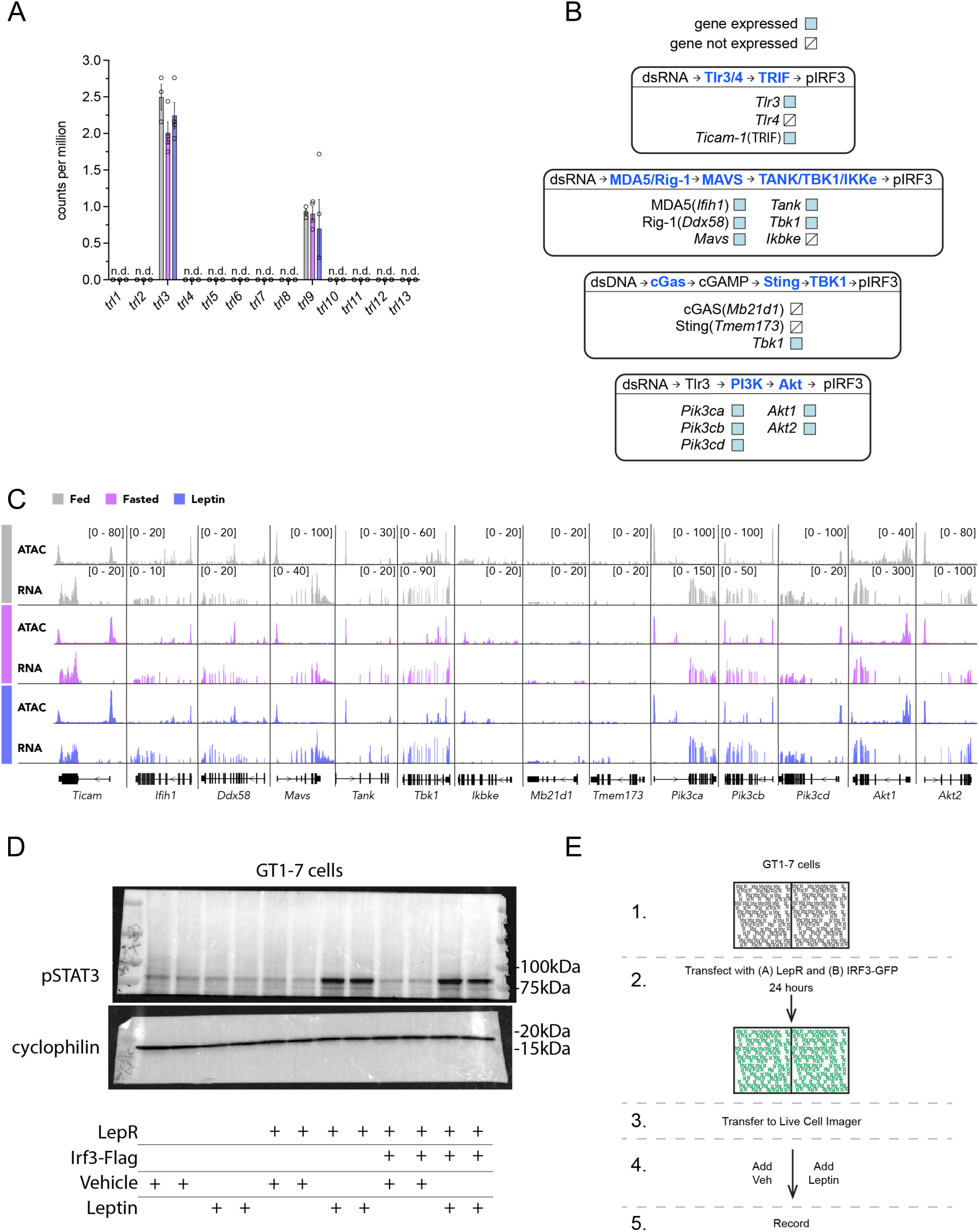
Establishment of a system for studying activation of IRF3 by leptin signaling. Associated with main Figure 4. **(A)** mRNA expression from NuTRAP^AgRP^ TRAP-seq experiment, showing the expression of all 13 toll-like receptors. Gene not detected (n.d.). **(B)** Relevant pathways for IRF3 activation, with blue boxes indicating the expression of the gene in our AgRP neuron RNA-seq dataset. **(C)** Genome browser views (IGV) showing AgRP neuron RNA-seq and ATAC-seq tracks for genes associated with IRF3 activation. **(D)** Western blot showing the leptin-responsivity of transfected GT1-7 cells treated with leptin. **(E)** Schematic of *in vitro* GT1-7 live-cell imaging experiment.

**Supplementary Figure 5.**
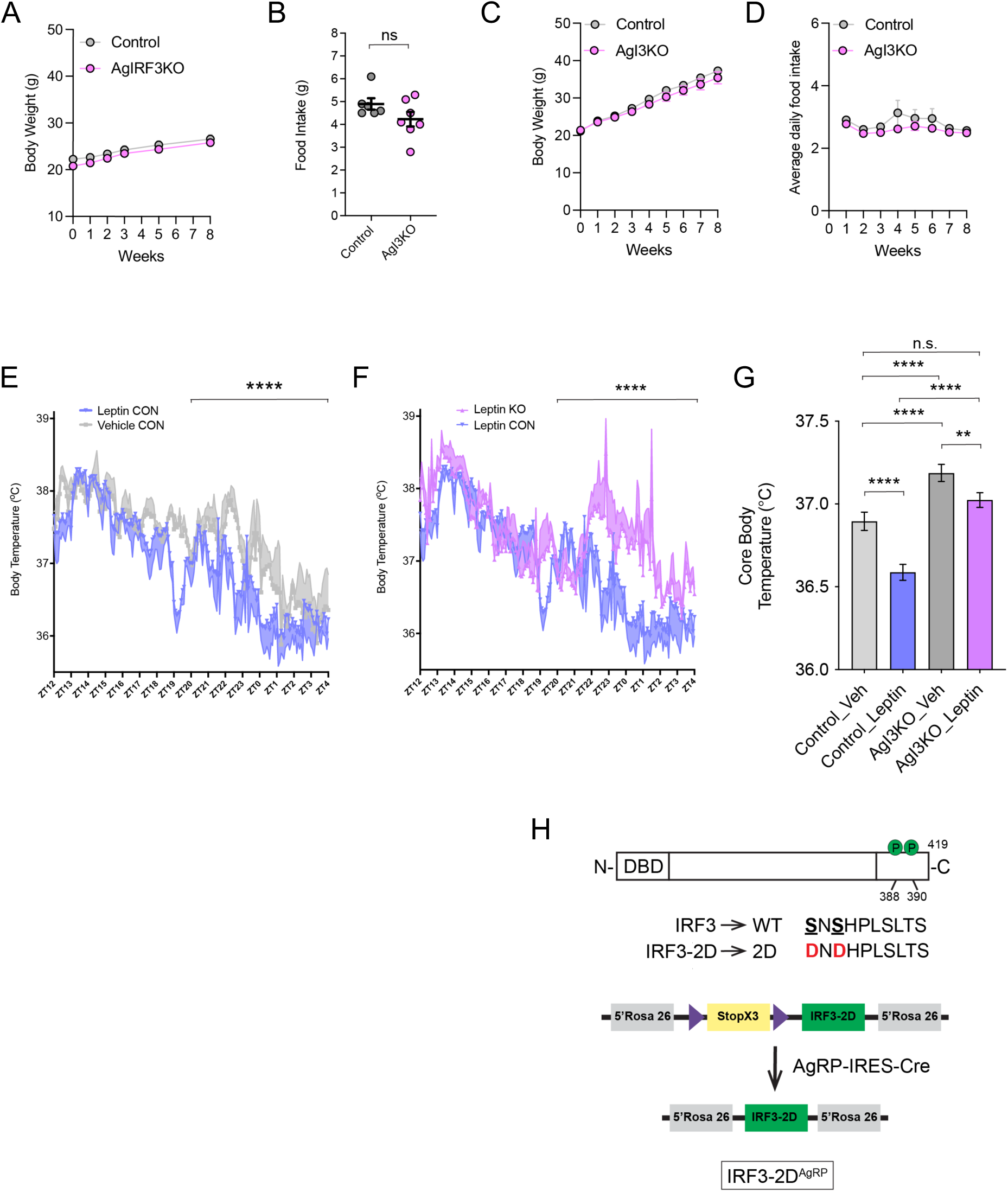
IRF3 loss-of-function and gain-of-function models in AgRP neurons. Associated with main Figure 5. **(A)** Body weight of chow-fed control and AgI3KO mice over an 8-week period. **(B)** 24-hour food-intake of chow-fed control and AgI3KO mice. **(C)** Body weight of high fat diet-fed control and AgI3KO mice over an 8-week period. **(D)** Weekly food-intake of high fat diet-fed control and AgI3KO mice. **(E)** Core body temperature measures during a 16-hour period showing control mice that received vehicle-t and leptin-treatment (5mg/kg). **(F)** Core body temperature measures during a 16-hour period following leptin-treatment (5mg/kg) in control mice or AgI3KO mice. Note that the leptin treated control cohort was the same for panels E and F, but the data are displayed separately to make distinctions between the different comparators clearer. **(G)** Quantification of core body temperature experiments. **(H)** Schematic of AgI3-2D mouse. Results are mean ± SEM of each condition and analyzed by a one-way ANOVA with repeated measures. *p < 0.05, **p < 0.01, ***p < 0.001, ****p < 0.0001.

**Supplementary Figure 6.**
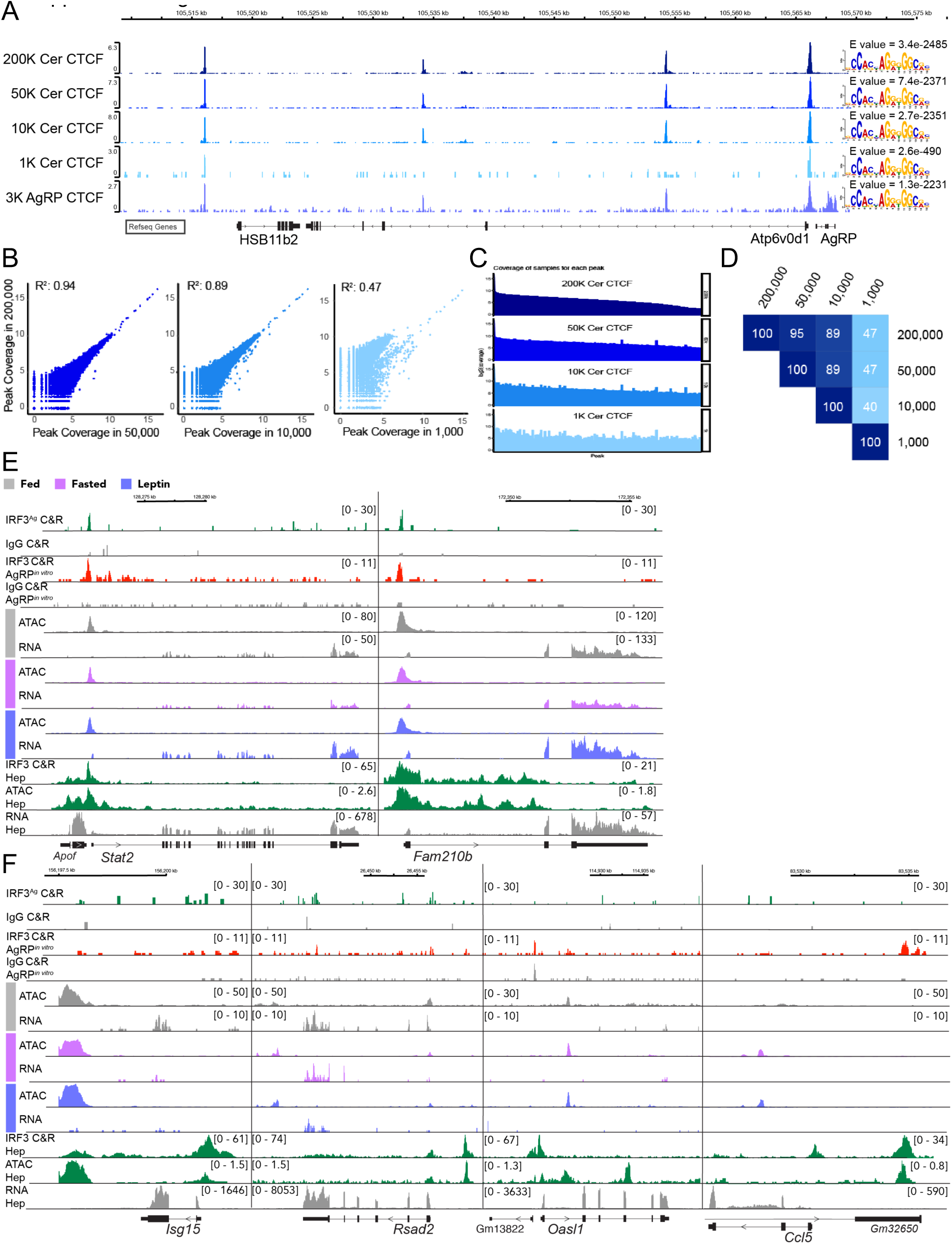
Determination of the CTCF cistrome in AgRP neurons. Associated with main Figure 6. **(A)** Genome browser views (IGV) of the locus containing *Hsb11b2*, *Atp6v0d1*, and *Agrp* are shown with whole-cerebellum CTCF CUT&RUN data titrated from 200,000 to 1,000 whole cerebellum nuclei, and a single 3,000 AgRP neuronal nuclei sample. CTCF motif e-values and position weight matrices are displayed for each condition. **(B)** Correlations of peak coverage between 200,000 nuclei (y axis) and each of the lower inputs (x axis) as indicated. Pearson correlation coefficients (R^2^) are shown in each plot. Each dot indicates coverage of an individual peak. **(C)** Peak coverage patterns in the titration. Coverage (log_2_) of peaks is shown as bars in an ascending order based on peak coverage in 200,000 nuclei. **(D)** Overlap of peaks in each pairwise comparison of the inputs. The number in each box indicates the percentage of the peaks overlapping in the pair set. **(E)** Top panel: Representative genes (*Stat2*, *Fam210b*) with a robust IRF3-2D^AGRP^ CUT&RUN peak near suspected leptin IRF3^AgRP^ target genes. For tracks from top-to-bottom: Representative *in vivo* IRF3-2D^AgRP^ IRF3 CUT&RUN, peri-hypothalamus IRF3-2D IgG CUT&RUN, *in vitro* immortalized AgRP neuron Adeno-IRF3-2D IRF3 CUT&RUN, *in vitro* immortalized AgRP neuron Adeno-IRF3-2D IgG CUT&RUN, NuTRAP^AgRP^ fed, fasted, leptin-treated ATAC-seq peaks, mouse hepatocyte primary culture Adeno-IRF3-2D IRF3 CUT&RUN, ATAC-seq and RNA-seq, all near the indicated genes. **(F)** Classic inflammatory IRF3 target genes not appreciably detected (*Isg15*, *Oasl1*, *Ccl5*), or very lowly detected (*Rsad2*) in NuTRAP^AgRP^ neurons, with corresponding, CUT&RUN, ATAC-seq, and RNA-seq tracks in the same order in the top panel.

**Table S1.**
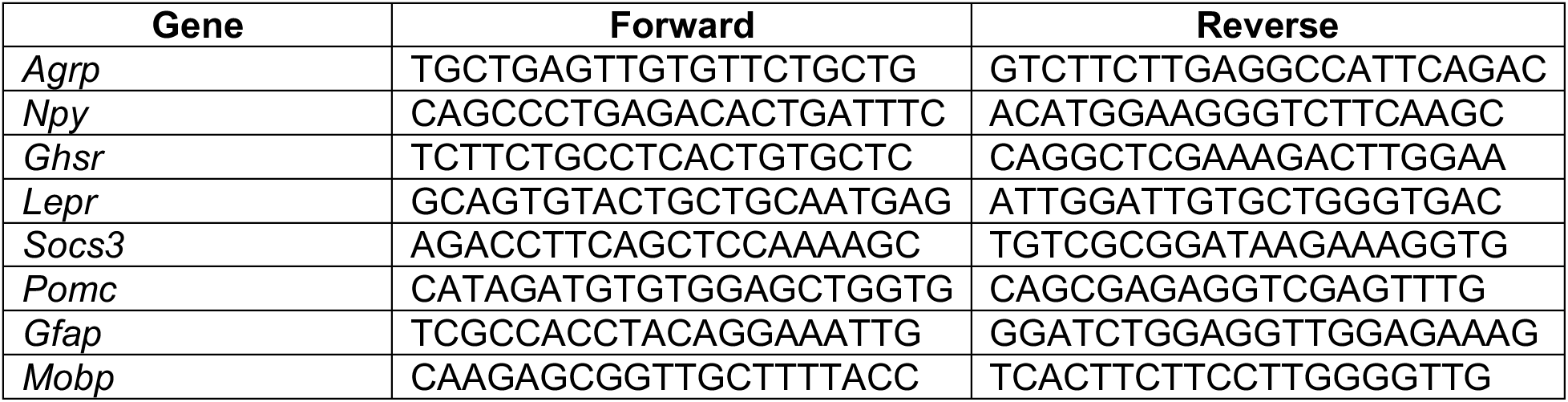
Primers used for qRT-PCR

